# Lysosomal alterations and decreased electrophysiological activity in CLN3 disease (966 bp deletion, E295K) patient-derived cortical neurons

**DOI:** 10.1101/2022.04.28.489465

**Authors:** Sueanne Chear, Sharn Perry, Richard Wilson, Aidan Bindoff, Jana Talbot, Tyson L Ware, Alexandra Grubman, James C Vickers, Alice Pébay, Jonathan B Ruddle, Anna E King, Alex W Hewitt, Anthony L Cook

## Abstract

CLN3 disease is a lysosomal storage disorder associated with fatal neurodegeneration that is caused by mutations in *CLN3*. Most individuals with CLN3 disease carry at least one allele with a 966 bp deletion in *CLN3* which results in the deletion of exons 7 and 8. There is a need for more physiologically relevant human cell-based CLN3 disease models to better understand the cellular changes during the disease process. Using CRISPR/Cas9, we corrected the 966 bp deletion mutation in human induced pluripotent stem cells (iPSCs) of a compound heterozygous patient (*CLN3* Δ 966 bp and E295K). The isogenic deletion-corrected and unedited CLN3 patient iPSCs were used for disease modeling. iPSC-derived neurons carrying this particular CLN3 mutation (CLN3 neurons) had lower functional activity as recorded using microelectrode arrays for most of the culture period. Proteomics analysis showed downregulation of proteins related to axon guidance and endocytosis at day *in vitro* (DIV) 14 and 42 in CLN3 neurons. This was accompanied by an increase in lysosomal-related proteins in CLN3 neurons. Western blot analysis revealed hyperglycosylation of the lysosomal marker, Lysosome Associated Membrane Protein 1 (LAMP1) in CLN3 neurons at DIV 14, 28 and 42, which was not apparent in control neurons. Ultrastructural analysis of CLN3 neurons showed numerous membrane-bound vacuoles containing diverse types of storage material, ranging from curvilinear deposits, multilamellar structures to osmiophilic deposits. Our findings suggest alterations in lysosomal function and neurodevelopment involving axon guidance and synaptic transmission in CLN3-deficient neuronal derivatives, which could be potential targets for therapy.

## INTRODUCTION

CLN3 disease (also known as Batten Disease or juvenile neuronal ceroid lipofuscinosis), is a lysosomal storage disorder caused by mutations in *CLN3*. There are more than 70 reported mutations of the *CLN3* gene causing frameshift with a premature stop codon, nonsense mutations, splice defects and missense mutations. The most prevalent mutation is a 966 bp deletion spanning exons 7 and 8, often referred to as a 1 kb deletion [22, 34], where approximately 75% of patients are homozygous for this mutation and ∼20% of patients are compound heterozygous for this and one of the other rare mutations [33].

*CLN3* encodes for a 438-amino acid protein, which appears to be a hydrophobic six-transmembrane domain protein [24]. The role of CLN3 in healthy and disease cells remains poorly understood, despite the identification of *CLN3* as the causative gene for CLN3 disease more than 20 years ago [34]. Studies from lower mammalian and other eukaryotic models have suggested that CLN3 is linked to endocytic trafficking [60], autophagy [6], cytoskeletal organization [51], osmoregulation [40], cell migration [20], cell viability [40], and lipid processing [13]. The hallmark of CLN3 disease is the formation of autofluorescent lipopigments with fingerprint profiles, which are enriched in subunit c of mitochondrial ATP synthase [14, 47], however, the underlying molecular basis behind subunit c accumulation remains unknown.

CLN3 disease typically presents early in children at between 4 and 8 years of age and is marked by vision failure, cognitive decline, behavioral changes, motor deficit, refractory seizures and eventually death [46, 54]. The neurological features are accompanied by reductions in whole brain and cerebral cortex volume mainly due to cortical and cerebellar atrophy and increased cerebrospinal fluid volume [2, 4]. Loss of Purkinje cells and granular cells in the cerebellum and dentate nucleus [1], neuron loss and glial activation in the hippocampus have also been described in CLN3 post-mortem brain tissue [68]. Characterization of cellular and molecular alterations can be provided by post-mortem tissue, however, these are usually limited to the late-stage pathological changes, are not amenable to intervention in the course of disease, nor provide longitudinal data. Mouse models are valuable in uncovering disease mechanisms and providing insights into the function of specific genes, especially in neurodegenerative disorders, however, the findings have rarely translated into human therapeutics [15].

For these reasons, human induced pluripotent stem cells (iPSCs) have emerged as a powerful tool for disease modeling and drug screening. When combined with CRISPR/Cas gene editing, patient-specific isogenic iPSCs can be generated and differentiated into neurons to model CLN3 disease, thus overcoming genetic variability between different donors as confounding factors [32]. More importantly, these patient-specific iPSC-derived models can provide insights into the precise roles of mutation and genetic background in the progression of disease. This is especially relevant in CLN3 disease, as even among CLN3 affected family members with identical mutations, variations in symptomatology are observed, possibly owing to genetic and environmental factors [11]. The differentiation of patient-specific iPSCs into neurons provides access to human neurons, one of the cell types relevant to neurological manifestations in CLN3 patients. Studies have demonstrated the relevance of iPSC-differentiated neurons harboring CLN3 homozygous and compound heterozygous 966 bp deletion in recapitulating certain CLN3 phenotypes such as fingerprint deposits and accumulation of subunit c of mitochondrial ATP synthase [30, 36]. In this study, isogenic CLN3 iPSCs derived from a compound heterozygous patient (*CLN3* Δ 966 bp and E295K) were differentiated into neurons to generate a robust disease model to investigate early-stage neuropathology. Our findings provide further insight into lysosomal and neurodevelopmental alterations caused by a CLN3 mutation.

## MATERIALS AND METHODS

### Human fibroblast culture

Primary fibroblasts, MBE 02873 (*CLN3* Δ 966 bp and E295K, hereafter designated CLN3 line) were established from a patient skin biopsy as described [44]. Fibroblasts were grown in medium containing 88% DMEM with high glucose (Gibco) supplemented with 10% fetal bovine serum (Gibco), 1% non-essential amino acids (Gibco) and 1% antibiotic-antimycotic (Gibco) in T75 flasks.

### Generation of iPSCs from CLN3 patient fibroblasts

CLN3 fibroblasts were reprogrammed using Epi5 Episomal iPSC reprogramming kit (Invitrogen). The episomal vectors encoding *OCT4, SOX2, LIN28, KLF4 and L-MY*C were transfected using Lipofectamine 3000 reagent (Invitrogen). Live surface marker staining with Anti-TRA-1-60 488 Live cell stain (Miltenyi Biotec) was performed at day 21 post-lipofection to identify undifferentiated iPSC colonies in the culture. Newly reprogrammed iPSCs were maintained in mTeSR plus medium (Stemcell Technologies) on Matrigel-coated plates.

### Assessment of iPSC pluripotency, differentiation potential and cell line identification

iPSCs were stained with primary antibodies rabbit anti-NANOG (1:200), rabbit anti-OCT4 (1:200), rabbit anti-PAX6 (1:200), mouse anti TRA-1-60 (1:200), mouse anti TRA-1-81 (1:200) and mouse anti-SSEA4 (1:200) (all from Cell Signaling Technology, StemLight Pluripotency Antibody Kit) to assess for pluripotency. To further assess pluripotency and trilineage differentiation potential of iPSC, embryoid bodies (EB) were formed. iPSCs at 80% confluency were dissociated with ReLeSR (Stemcell Technologies). Cell aggregates were transferred into ultra-low attachment 6-well plates in EB medium consisting of 20% knockout serum replacement (Gibco), 79% DMEM/F-12, GlutaMAX supplement (Gibco), 1% non-essential amino acids and 0.1 mM 2-mercaptoethanol (Sigma) for EB formation. Media was refreshed every 2 days. On day 4, EBs were plated onto Matrigel-coated wells for further differentiation in EB medium for 10 days. Expression of pluripotency and germ layer genes of EBs on day 14 of differentiation was examined using TaqMan hPSC Scorecard Panel (Thermo Fisher Scientifc) following manufacturer’s instructions. Data analysis was done using web-based hPSC scorecard analysis software. Short tandem repeat (STR) profiling and analysis of CLN3 fibroblasts and iPSCs were carried out by Australian Genome Research Facility (Melbourne, Australia).

### Detection of episomal vector

To detect the presence of episomal vectors in reprogrammed iPSCs, endpoint PCR was performed on genomic DNA extracted from the iPSCs using EBNA-1 primers (Additional file 1: Table S1) which identifies all five episomal plasmids in the Epi5 reprogramming kit.

### Virtual karyotyping analysis

Copy number variation (CNV) analysis of parental fibroblasts and iPSCs was performed using Illumina HumanCytoSNP-12 beadchip array (Illumina). B allele frequency and log R ratio of each single nucleotide polymorphism (SNP) marker were collected from GenomeStudio (Illumina) and analyzed with PennCNV [70] and QuantiSNP [10] with default parameter settings. Genomic regions with CNV calls having at least 20 contiguous SNPs or genomic regions with SNPs spanning at least 1 Mb generated by both PennCNV and QuantiSNP were then visualized using GenomeStudio to confirm the absence of chromosomal aberration [44].

### Design of crRNA and donor plasmid

crRNA, CAAGGTAGGGACTTGAAGGA, which targets the breakpoint sequence of CLN3 966 bp deletion, was designed using the Benchling platform (www.benchling.com). A ∼3.7 kbp donor repair construct was cloned into pUC57 vector at the EcoRV site by GenScript (Nanjing, China). The donor plasmid consists of the corrected *CLN3* sequence with loxP-flanked puromycin-selection cassette flanked by 801-911 bp of homologous sequence.

### CRISPR/Cas9-mediated correction of 966 bp deletion

Editing of CLN3 iPSCs to correct 966 bp deletion was done essentially as described [71]. Single cell suspension (800,000 cells) of CLN3 iPSCs was electroporated with Cas9-sgRNA ribonucleoprotein (IDT) and 966 bp donor plasmid using Amaxa 4D Nucleofector (Lonza) with program CB 150. iPSCs were treated with puromycin 72 hours after CRISPR/Cas9 electroporation when cells have reached ∼80% confluency. Cells were cultured in 0.125 µg/mL (half the kill dose) puromycin in mTeSR plus medium for 3 days, with daily media replacement. Cells were then cultured in 0.25 µg/mL puromycin in mTeSR plus medium for another 3 days. When distinct colonies appeared large enough for picking (∼200 µm), they were isolated for further expansion and genotyping. DNA extract from edited clones were amplified through PCR reaction using CLN3 F1 and CLN3 R1 primers that span the 966 bp deletion region. Successfully-edited cells, as confirmed by PCR were treated with Cre recombinase gesicles (Takara Bio) with 6 µg/mL Polybrene (Sigma) in mTeSR plus medium to remove the puromycin-resistance cassette. Successful Cre-Lox recombination was confirmed by PCR using CLN3 F1 and R1 primers. Sanger sequencing of cDNA of the edited clone was performed using primers CLN3_SS_cdna_F1 and CLN3_SS_cdna_R1. All primers used are listed in Additional file 1: Table S1.

### Off-target analysis

The top 10 predicted off-target sites and all predicted off-target sites within exonic regions were determined using Benchling (www.benchling.com). Sanger sequencing was performed to sequence the 300-400 bp sequences surrounding the off-target sites in both edited and native unedited cell lines. Indels were analyzed with Synthego ICE analysis tool (https://ice.synthego.com/#/).

### Neural differentiation

Neural induction and differentiation were done essentially as described [65]. When the iPSCs were 80% confluent, cells were dissociated into small clumps using Accutase (Gibco) and seeded at a density of 3 × 10^5^ cells/well in poly-l-ornithine (PLO, Sigma)/laminin (Gibco)-coated 6-well plate. Cells were cultured in mTeSR plus medium supplemented with 10 µM Rho-associated protein kinase (ROCK) inhibitor (Stemcell Technologies). After overnight incubation, the spent media was replaced by Neural Induction Medium containing 98% Neurobasal medium (Gibco) and 2% neural induction supplement (Gibco) which was subsequently refreshed every other day. On day 7 of culture, primitive neural stem cells (NSCs) were dissociated with Accutase and seeded at a density of 8 × 10^5^ cells/well in a PLO/laminin-coated plate. Cells were cultured in Neural Expansion Medium (NEM) consisting of 49% Neurobasal medium, 49% Advanced DMEM (Gibco) and 2% neural induction supplement pre-treated with 5 µM ROCK inhibitor. NEM without ROCK inhibitor was changed every other day until cells were confluent.

For differentiation into mature neurons, NSCs at passage 4 were plated onto PLO/laminin-coated 6-well plates. When cells were 75-90% confluent, NEM was replaced with neuronal differentiation medium consisting of 95% Neurobasal plus medium (Gibco) supplemented with 2% B-27 Plus (Gibco), 1% GlutaMAX (Gibco), 1% CultureOne (Gibco) and 1% antibiotic-antimycotic. The next day, half-media exchange was performed. On day 4, cells were dissociated with Accutase and seeded into PLO/laminin-coated 6-well plates containing neuronal maturation medium (95% Neurobasal plus, 2% B-27 Plus, 1% GlutaMAX, 1% CultureOne, 1% antibiotic-antimycotic supplemented with 10 ng/mL BDNF (StemCell Technologies), 10 ng/mL GDNF (StemCell Technologies), 0.25 mM db-cAMP (Sigma) and 200 µM L-Ascorbic acid (Sigma). Cells were cultured for 42 days in neuronal maturation medium. Media was replaced with fresh prewarmed media every 2-3 days with half volume change.

### RNA extraction and RT-qPCR

RNA was extracted from iPSCs, NSCs, and neurons at day *in vitro* (DIV) 7, 14, 28 and 42 using the RNAeasy Plus Mini Kit (Qiagen). cDNA was synthesized using 1 µg of RNA per sample according to manufacturer’s instructions (Omniscript RT kit, Qiagen). Taqman assay probes (Thermo Fisher Scientific) used for RT-qPCR are listed in Additional file 1: Table S2.

### Microelectrode array

NSCs from CLN3 and CLN3-Cor were differentiated into neurons in parallel (see neural differentiation methods). On differentiation day 4, 150,000 cells were replated as a 75 µL drop directly onto the PLO/laminin coated electrode array of each well of 24-well microelectrode array (MEA) plates (MultiChannel Systems, Germany), where each well was fitted with 12 PEDOT-coated gold electrodes. Eight wells per independent culture were dedicated to each genotype (n=3 independent cultures). After 30 minutes, 500 µL of neuronal maturation medium was added gently into each well. Cells were incubated at 37 °C, 5% CO2 for 4 days before the first MEA recording. Half media exchanges were performed every 2-3 days. On experimental days, the MEA plate was removed from the incubator, placed on the MEA heated stage (MultiChannel Systems, Germany) and maintained at 37 °C throughout the duration of data acquisition. Plates were left to equilibrate for 2 minutes prior to starting recordings. Media exchanges were performed after recording. Neuronal activity from each well was recorded with the MultiWell-MEA-system (MultiChannel Systems, Germany). MEA recordings were acquired in Multiwell-Screen Version 1.11.7.0, low pass filtered at 3500 Hz, high pass filtered at 1 Hz, and sampled/digitized at 20 kHz. Spontaneous network activity was recorded for 5 minute intervals, once per day, starting from DIV 4 to 42. Neuronal activity was analyzed using MultiWell-Analyzer Version 1.8.7 and a customized R script. Wells containing electrodes that detected ≥ 10 spikes/min with minimum amplitude of 20 µV were considered active wells. Wells with at least 1 active electrode were included in analysis. The adaptive spike detection threshold for each electrode was set at 5 times the standard deviation of the baseline noise level during the first second of reading for each electrode with 1 s binning. Burst activity was detected using the following threshold parameters: maximum interval to start burst, 50 ms; maximum interval to end burst, 50 ms; minimum interval between bursts, 100 ms; minimum duration of burst, 50 ms; and minimum spike count in burst; 4 (Additional file 2: Figure S1a). Network bursts were defined as synchronous network spikes and bursts in which three or more electrodes (out of a total of 12) capture activity simultaneously. MEA parameters and their definitions are shown in Additional file 1: Table S3. The values from each active electrode in each active well were averaged to determine a ‘well average’ for further analysis.

### Protein preparation for mass spectrometry

Proteins were harvested from CLN3 and CLN3-Cor neurons (n=3 independent cultures/cell line) which had been plated in 12-well plates. Protein was extracted using 100 µL of lysis buffer (7 M urea (Sigma), 2 M thiourea (Sigma) and 30 mM Trizma base (Sigma) containing protease (Roche) and phosphatase inhibitors (Roche)). Cell lysates were sonicated for 3 cycles of 15 seconds pulse with 5 seconds interval on ice. Cell lysates were mixed gently on a rotary suspension mixer at 4 °C for 2 hours, centrifuged at 13,000 rpm for 15 minutes before supernatant was collected. Protein concentration was determined by performing Pierce™ 660 nm Protein Assay (Thermo Fisher Scientific) according to manufacturer’s instructions. Proteins (30 µg/sample) were sequentially reduced and alkylated then cleaned up using the SP3 method [25], followed by digestion with 1.2 µg proteomics grade trypsin/rLysC (Promega) overnight at 37 °C.

### Mass spectrometry – data-independent acquisition (DIA)

Peptide samples were analyzed by nanoflow high-performance liquid chromatography-mass spectrometry/mass spectrometry (MS) using an Ultimate 3000 nano RSLC system (Thermo Fisher) coupled with a Q-Exactive HF mass spectrometer fitted with a nanospray Flex ion source (Thermo Fisher) and controlled using Xcalibur software (version 4.3). Approximately 1 µg of each sample was injected and separated using a 120-minute segmented gradient by preconcentration onto a 20 mm x 75 µm PepMap 100 C18 trapping column then separation on a 250 mm x 75 µm PepMap 100 C18 analytical column at a flow rate of 300 nL/min and held at 45 °C. MS Tune software (version 2.9) parameters used for data acquisition were: 2.0 kV spray voltage, S-lens RF level of 60 and heated capillary set to 250 °C. MS1 spectra (390 – 1240 m/z) were acquired at a scan resolution of 120,000 in profile mode with an AGC target of 3e6 and followed by sequential MS2 scans across 26 DIA x 25 amu windows over the range of 397.5-1027.5 m/z, with 1 amu overlap between sequential windows. MS2 spectra were acquired in centroid mode at a resolution of 30,000 using an AGC target of 1e6, maximum IT of 55 ms and normalized collision energy of 27.

### Mass spectrometry raw data processing and statistical analysis

DIA-MS raw files were processed using Spectronaut software (version 14.8, Biognosys AB). A project-specific library was generated using the Pulsar search engine to search the DIA MS2 spectra against the *Homo sapiens* UniProt reference proteome concatenated with common contaminants (comprising 20,455 entries, September 2018). With the exception that single-hit proteins were excluded, default (BGS factory) settings were used for both spectral library generation and DIA data extraction. For library generation, these included N-terminal acetylation and methionine oxidation as variable modifications and cysteine carbamidomethylation as a fixed modification, up to two missed cleavages allowed and peptide, protein and PSM thresholds set to 0.01. Mass tolerances were based on first pass calibration and extensive calibration for the calibration and main searches, respectively, with correction factors set to 1 at the MS1 and MS2 levels. Targeted searching of the library based on XIC extraction deployed dynamic retention time alignment with a correction factor of 1. Quantitation was based on the MaxLFQ algorithm for derivation of inter-run peptide ratios, followed by cross-run normalization based on median peptide intensity.

Differential abundance of proteins between genotypes at different time points was determined using One-way ANOVA followed by post-hoc test with Student’s T-test with a permutation-based false discovery rate (FDR) set at 0.05 and S0=0.1 to exclude proteins with very small difference between means (Perseus software version 1.6.14.0). Principal component analysis (PCA) plot of the isogenic CLN3 neuronal proteome was generated in Spektronaut software. UNIPROT accessions for proteins that were significantly increased or decreased in abundance in CLN3 neurons compared to CLN3-Cor neurons were uploaded to DAVID bioinformatics resource 6.8 to retrieve the enriched Kyoto Encyclopedia of Genes and Genomes (KEGG) pathways. KEGG pathways with *p* < 0.05 after Benjamini-Hochberg correction were considered significant. The mass spectrometry proteomics data for this project have been deposited to the ProteomeXchange Consortium via the PRIDE [55] partner repository with the dataset identifier PXD032191.

### Western blotting

Proteins from neuronal culture at DIV 14, 28 and 42 grown in a 12-well plate were extracted using 50 µL ice-cold RIPA lysis buffer (Abcam) added with protease inhibitor (Roche) and phosphatase inhibitor (Roche). Protein concentration was determined by performing Pierce BCA Protein assay according to manufacturer’s instructions (Thermo Fisher Scientific). After Western blotting, membranes were incubated in LAMP1 (Cell Signaling, D2D11; 1:4000), subunit c ATP synthase (Abcam, ab181243; 1:4000) and the loading control, GAPDH (Merck, AB2302, 1:120,000). Densitometric analysis of protein bands was performed using Image Studio Lite Version 5.2 to derive band intensities which were then normalized to GAPDH.

### Deglycosylation of LAMP1

Proteins from neuronal culture at DIV 14 were extracted as described above and treated with and without N-glycanase or endoglycosidase H according to manufacturer’s instructions (New England Biolabs). The lysates were analyzed with Western blot using LAMP1 (1:4000) antibody.

### Transmission electron microscopy

Neuronal culture in confluent 12-well plates were fixed directly with gentle agitation for 15 minutes at room temperature with a fixative containing 2.5% glutaraldehyde (Electron Microscopy Sciences), 2% paraformaldehyde (PFA, Electron Microscopy Sciences), 0.025% calcium chloride (Sigma) in a 0.1 M sodium cacodylate (Electron Microscopy Sciences) buffer pH 7.4. Cells were washed (3 × 10 minutes) with cacodylate buffer. Cells were scraped gently into a microfuge tube and postfixed in 1% osmium tetroxide (Electron Microscopy Sciences) for 30 minutes - 1 hour at 4 °C. Cells were washed with Milli-Q water (3 × 10 minutes), followed by overnight staining with 0.5% uranyl acetate (Electron Microscopy Sciences) at 4 °C. The next day, uranyl acetate was activated through heating at 50 °C for 1 hour. Cells were washed with Milli-Q water (3 × 10 minutes) and dehydrated in graded ethanol solutions and propylene oxide on ice in the following order: 50%, 70%, 80%, 90%, 100% (5 minutes each), propylene oxide (3 × 5 minutes) and infiltrated overnight with propylene oxide: resin (1:1) overnight. The next day, cells were embedded in resin and polymerized for two days at 60 °C. After polymerization, thin sections (70 nm) from resin blocks were cut and stained with Uranyless solution (ProSciTech) for 7 minutes followed by lead citrate (Sigma) for 7 minutes. Images were captured with Hitachi 7700 transmission electron microscope with a LaB6 filament, at 80 kV in high contrast mode.

The area of late degradative autophagic vacuoles having one limiting membrane with cytoplasmic materials [22] was quantitated in 20-21 cells from 3 independent batches per genotype. Regions of interest were outlined using ImageJ to quantitate the autophagic vacuole area.

### DQ-Bovine serum albumin assay

Cells were seeded at a density of 25,000 cells/well in a black flat bottom tissue culture treated 96-well plate. On the day of the experiment, cells were rinsed once with prewarmed DPBS^-/-^ (Gibco) then incubated with 10 µg/mL DQ Green Bovine Serum albumin (BSA, Invitrogen), which was diluted in medium for 6 hours at 37 °C, 5% CO2. After incubation, DQ Green BSA was removed, and cells were fixed with 4% PFA for 20 minutes at room temperature. Cells were washed with PBS (3 × 10 minutes) and incubated with 5 µg/mL DAPI for 10 minutes. Cells were imaged with Celldiscoverer 7 using At495 (495/527 nm) and DAPI channels. DQ-BSA puncta (diameter: 3-9 pixels) in cell bodies were segmented and area of DQ-BSA puncta, normalized to cell area, was quantified. Image analysis was performed using CellProfiler 4.2.1. Number of cells quantified were: DIV 14 (CLN3: 1584 cells, CLN3-Cor: 9266 cells), DIV 28 (CLN3: 1816 cells, CLN3-Cor: 7073 cells), DIV 42 (CLN3: 2033 cells, CLN3-Cor: 10323 cells). We estimated the mean DQ-BSA puncta area normalized to cell area using mixed-effects beta regression with the glmmTMB [5] package for R [56]. Beta regression was appropriate because the dependent variable was a proportion and thus bounded by 0 and 1, so in order to estimate confidence intervals with the appropriate coverage and bounding we assumed the residuals were beta-distributed. The model was fitted on the logit scale. Random intercepts were fitted to account for clustering within the experimental unit of batch. Inspection of residuals suggested that an excess of zeros was associated with time, so we tested a zero-inflated beta regression model using a likelihood ratio test, conditioning the zero process on time, which was significant.

### Statistical analyses

All statistical analyses, unless otherwise indicated, were performed using R [56] and *p* < 0.05 were considered significant. Data from MEA experiment were analyzed using a generalized additive model in the mgcv package [72]. For each MEA feature, we smoothed the response over time (DIV) by fitting a cubic spline with a shrinkage penalty. Data from western blots, electron microscopy and RT-qPCR studies were modeled using linear mixed effects models in the lme4 [3] package. To account for pseudoreplication, we fitted random intercepts (assumed to be normally distributed) for the clustering on batch. We used Tukey’s method with a correction for multiple comparisons to estimate differences between cell lines.

## RESULTS

### Generation of patient-specific CLN3 iPSCs

We reprogrammed fibroblasts from a CLN3 patient to pluripotency under feeder-free conditions using episomal vectors. A series of quality control tests was performed on the reprogrammed iPSC line. CLN3 iPSCs were integration-free by passage 7 as demonstrated by PCR on DNA using EBNA1-specific primers (Additional file 3: Figure S2a). Immunocytochemistry (Additional file 3: Figure S2b) results showed that at the protein level, CLN3 iPSCs expressed pluripotent markers. The Taqman Scorecard analysis demonstrated that the expression levels of pluripotency genes in iPSCs were comparable to undifferentiated embryonic stem cells whereas the iPSC-differentiated embryoid bodies displayed an expression pattern of trilineage genes comparable to germline-specific controls (Additional file 3: Figure S2c). Karyotyping of reprogrammed iPSCs revealed a normal male karyotype (46, X,Y) (Additional file 4: Figure S3). STR analysis of 10 different genomic loci indicated that the CLN3 iPSC line was 100% matched to the donor fibroblast DNA.

### CRISPR/Cas9-mediated correction of 966 bp deletion in CLN3 gene

To correct the 966 bp deletion mutation, we electroporated CLN3 iPSCs with an allele-specific single guide RNA (sgRNA) which targets the breakpoint sequence of the 966 bp deletion allele together with a homology-directed repair (HDR) donor plasmid carrying the reference 966 bp sequence and a floxed puromycin resistance gene for selection (Fig. 1a). Puromycin-resistant clones were validated for HDR correction via PCR using primer sets which target the region between intron 6 and intron 8 of *CLN3*. From PCR gel analysis, 966 bp correction was evident by larger size of amplicon ∼2.5 kbp which includes the puromycin sequence and the absence of the truncated 966 bp deletion band (Fig. 1b). To avoid alteration in the endogenous DNA sequence after HDR correction of CLN3 iPSCs, the corrected clone was treated with Cre recombinase to excise the puromycin casette. Following Cre-Lox recombination, the corrected clone demonstrated a single wild-type CLN3 band similar to a control cell line, along with absence of the puromycin band on PCR (Fig. 1b). PCR of cDNA from the corrected clone also showed a homozygous CLN3 band similar to that of a wild-type control (Fig. 1c). Sanger sequencing of the corrected clone cDNA showed restoration of exons 7 and 8 (Fig. 1d). The corrected cell line is hereafter referred to as CLN3-Cor. The CLN3-Cor iPSCs stained positive for pluripotency markers OCT4, SOX2, NANOG, TRA-1-60, TRA-1-81 and SSEA-4 (Additional file 5: Figure S4a). CLN3-Cor iPSC-derived embryoid bodies were able to express markers of all three germ layers, mesoderm, ectoderm and endoderm as demonstrated by Taqman hPSC scorecard assay (Additional file 5: Figure S4b). Virtual karyotyping analysis from both QuantiSNP and PennCNV methods showed normal male karyotype for CLN3-Cor iPSCs (46, XY) without CRISPR/Cas9 editing-induced chromosomal abnormalities (Additional file 6: Figure S5). Results from Synthego Ice Analysis Tool (Additional file 1: Table S4) did not reveal any sgRNA-induced modifications on the predicted off-target sites.

**Fig. 1.**
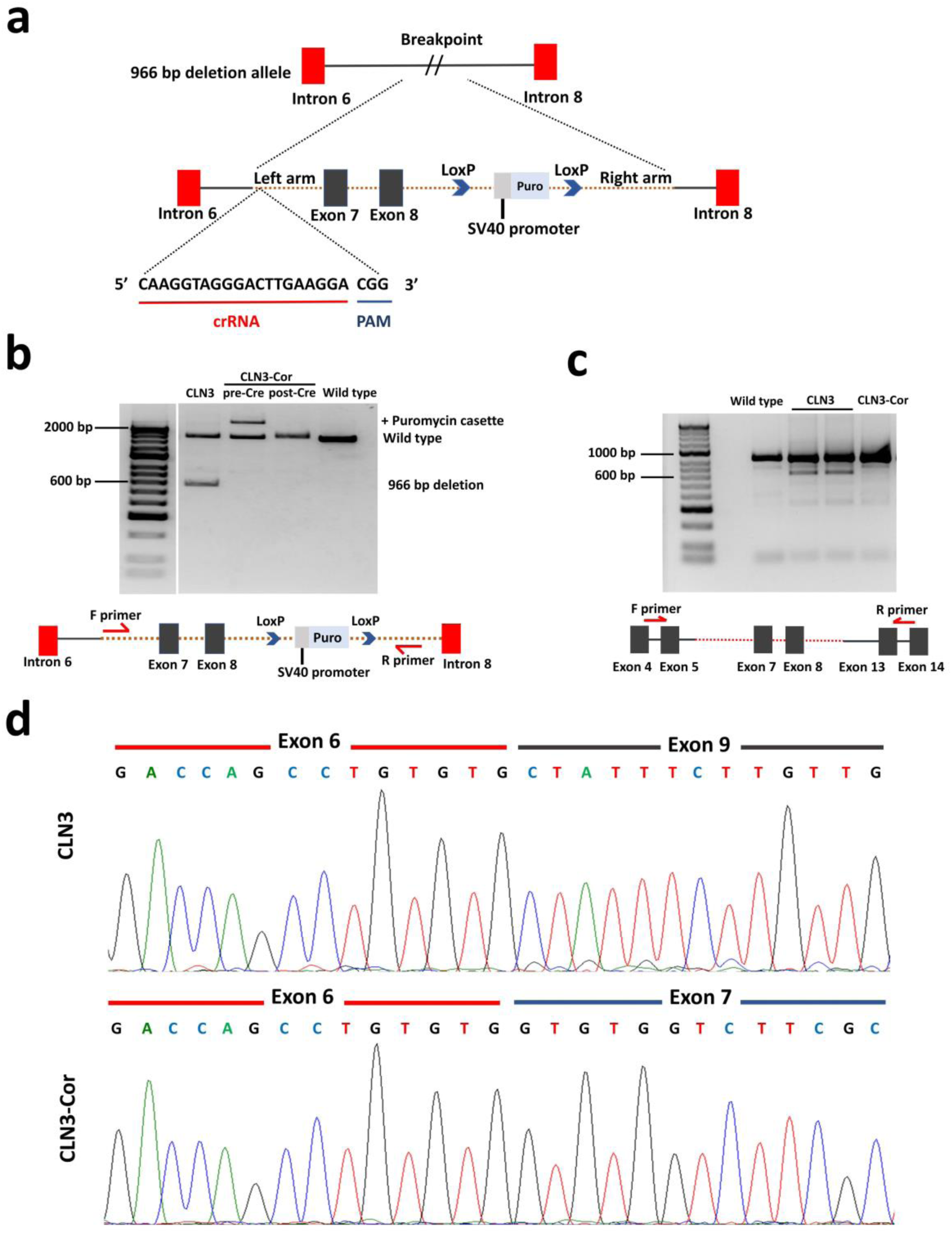
CRISPR/Cas9-mediated insertion of 966 bp sequence. (a) Schematic representation depicting donor repair construct carrying the reference CLN3 sequence and a loxP-flanked puromycin resistance cassette targeted to breakpoint sequence by crRNA. (b) Genomic PCR products of CLN3 iPSCs, CLN3-Cor iPSCs before and after Cre-Lox recombination and a wild-type positive control amplified using primers that span exons 7 and 8 of the CLN3 gene. (c) RT-PCR of wild-type positive control, CLN3 iPSCs and CLN3-Cor iPSCs. Diagrams below gel image depict location of primers. (d) Sanger sequencing of target cDNA regions revealed restoration of the deleted exons.

### CLN3 mutation does not affect differentiation efficiency of CLN3-deficient neurons

We generated human cortical neuron cultures from isogenic CLN3 iPSCs by inducing NSCs that expressed PAX6 and NESTIN (Fig. 2a). The neuronal culture stained positive for mature neuronal markers, MAP2 and TAU (Fig. 2b). RT-qPCR on RNA extracts from NSCs for both cell lines showed high expression of *NESTIN* and *PAX6* which then was reduced following neural differentiation (Additional file 7, Figure S6). Both isogenic neuronal cultures showed expression of immature (*TUBB3*) and mature (*MAP2*) neuronal markers, glutamatergic neuron-related genes (*GRIA2, GRIN1, SLC1A7, SLC1A2* and *SLC1A3*), GABAergic neuron-related genes (*GAD1, GAD2*) and postsynaptic marker (*DLG4*) (Additional file 7, Figure S6). At DIV 7, the expression of *GAD1, GAD, GRIA2, GRIN1, SLC1A2* and *DLG4* was higher in CLN3 than in CLN3-Cor neurons, while the expression of *TUBB3* and *SLC1A3* was lower in CLN3 than in CLN3-Cor neurons (Additional file 7, Figure S6). At DIV 28, the expression of *MAP2, GAD2, GRIA2, GRIN1, SLC1A2, DLG4* and *TUBB3* was lower in CLN3 compared with CLN3-Cor neurons (Additional file 7, Figure S6). Expression of *CLN3* increased when both isogenic CLN3 iPSCs were differentiated into neurons from NSCs. The CLN3-Cor cell line had higher expression of *CLN3* at several stages, including iPSCs, NSCs and neurons at DIV 28 and 42 (Additional file 7, Figure S6).

**Fig. 2.**
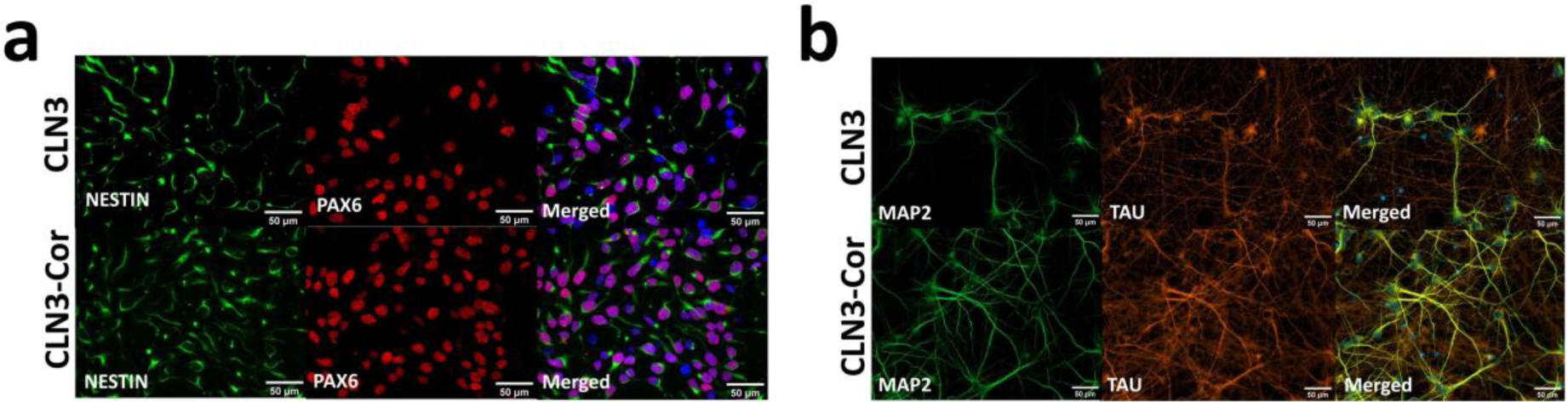
Neural differentiation of isogenic CLN3 iPSCs. (a) CLN3 and CLN3-Cor NSCs stained positive for NSC markers, NESTIN and PAX6 (b) CLN3 and CLN3-Cor neurons at DIV 28 expressed mature neuronal markers, MAP2 and TAU. Scale bars: 50 µm.

### Development of spontaneous firing activity in isogenic CLN3 cortical networks

Neurological abnormalities, for example cognitive decline, motor imbalance and refractory seizure, are some of the main presenting symptoms of CLN3 patients. To investigate if alterations in neuronal activity patterns may be present in neurons with CLN3 mutation, the electrophysiological activity of iPSC-derived CLN3 and CLN3-Cor neurons was recorded using an MEA, once a day from DIV 4 to DIV 42. Dissociated pre-maturation neurons started growing new neurites immediately after seeding followed by formation of interconnected circuits where both cultures displayed an even cell distribution on the MEA plates (Fig. 3a) and fired spikes, bursts and network bursts (Additional file 2, Figure S1b) throughout the culture period (Fig. 3b).

**Fig. 3.**
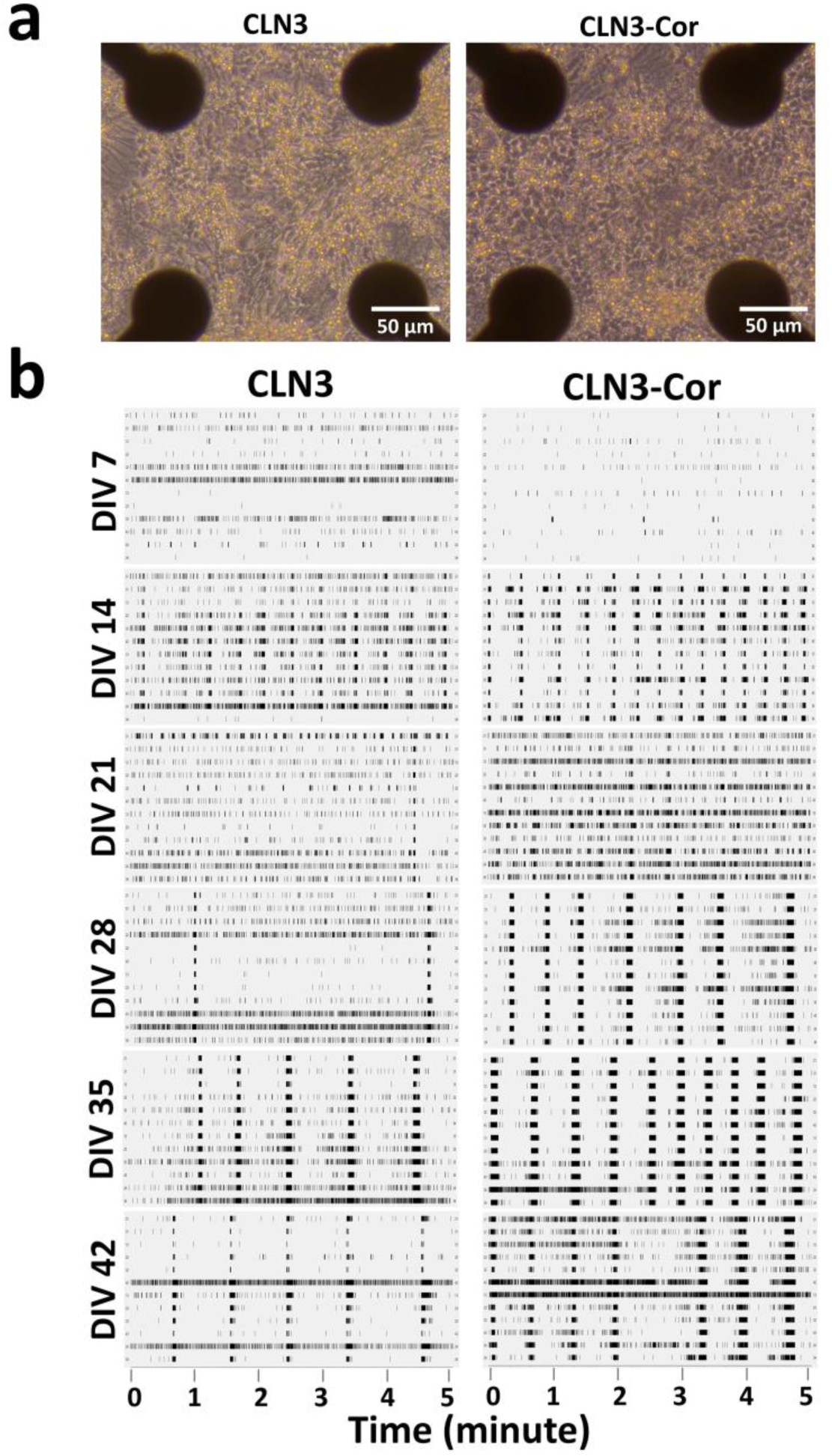
Electrophysiological activity of isogenic CLN3 neurons on MEA. (a) Representative images of CLN3 and CLN3-Cor neuronal cultures on MEA plates at DIV 10. (b) Representative raster plots of isogenic CLN3 neurons showing 5 minutes of electrophysiological activity across development from DIV 7 to DIV 42 (n=3 independent experiments; 8 wells/cell line/experiment)

The percentage of electrodes detecting activity (active electrodes) in both CLN3 and CLN3-Cor neurons peaked at DIV 32 (62.6%) and DIV 37 (97.1%), respectively (Fig. 4a). During DIV 4-8, the percentage of active electrodes was slightly higher in CLN3 neurons than in the CLN3-Cor neurons (mean CLN3: 20.8%; CLN3-Cor: 15.04%), however, from DIV 12 onward, CLN3 neurons had a lower percentage of active electrodes than CLN3-neurons (mean CLN3: 48.97%; CLN3-Cor: 81.27%; Fig. 4a). The spike rate of CLN3 neurons increased from DIV 4 onward, peaked at DIV 12 (2.27 Hz) and then remained consistent toward DIV 42 whilst in CLN3-Cor neurons, spike rate increased from DIV 4 and peaked at DIV 25 (15.02 Hz), before declining toward DIV 42 (4.37 Hz) (Fig. 4b). CLN3 neurons from DIV 4-10, had a higher spike rate than CLN3-Cor neurons (mean CLN3: 1.37 Hz; CLN3-Cor: 0.56 Hz), however, from DIV 12 onward, CLN3 neurons fired less frequently than CLN3-Cor neurons (mean CLN3: 2.36 Hz; CLN3-Cor: 10.53 Hz; Fig. 4b). In association with an increased spike rate, the interspike interval (ISI), the period of inactivity between spikes, in CLN3 neurons decreased between DIV 4-12 (DIV 4: 3.02 s; DIV 12: 1.24 s) before plateauing toward DIV 42 (Fig. 4c). A similar trend was seen in CLN3-Cor neurons where the ISI declined rapidly from DIV 4 (5.85 s) to DIV 17 (0.21 s) before plateauing toward the end of the culture period (Fig. 4c). The higher spike rate combined with the reduced ISI from DIV 12, indicates CLN3-Cor neurons are more functionally active than CLN3 neurons.

**Fig. 4.**
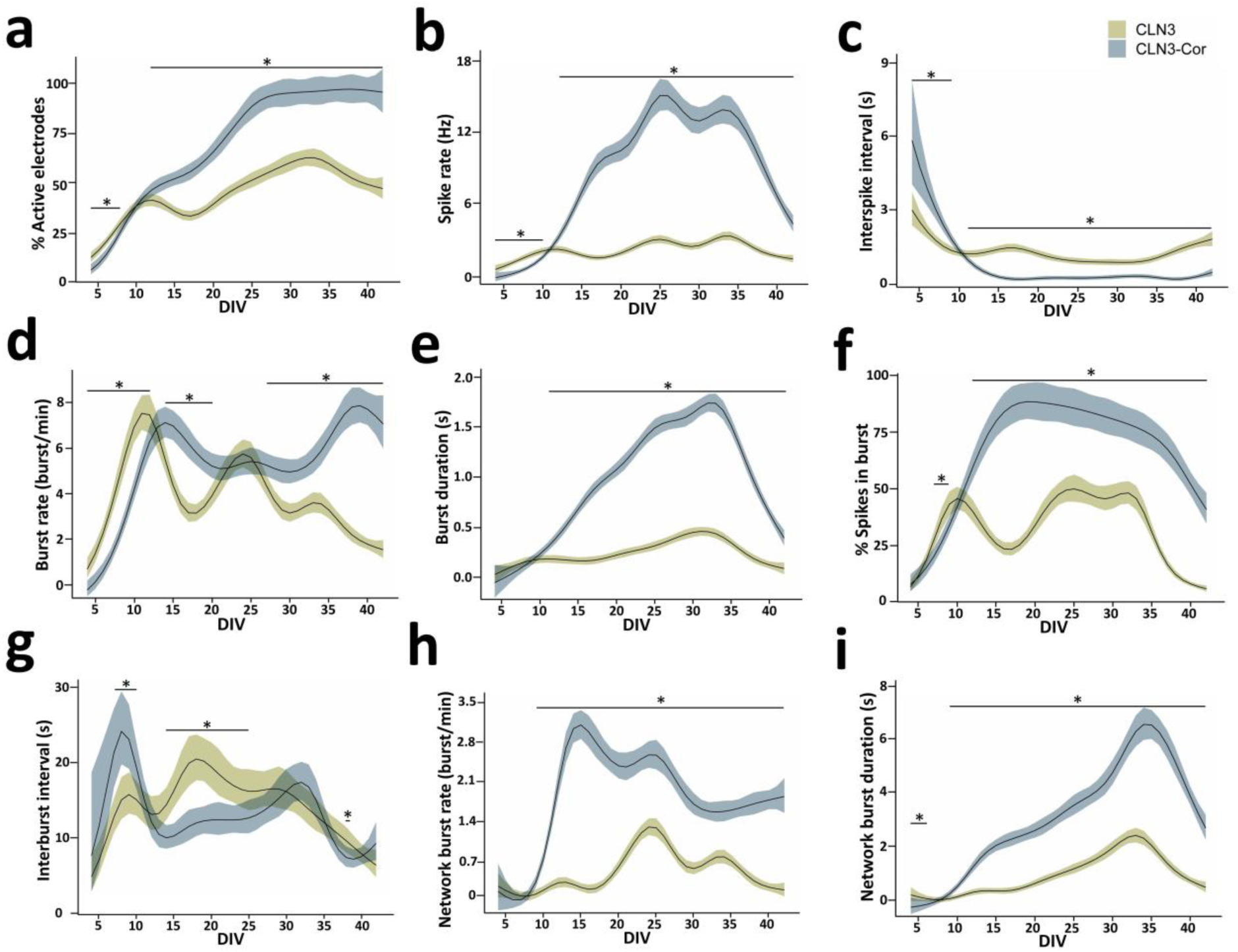
Development of functional activity of isogenic CLN3 neurons on MEA. Graphs showing development of (a-c) spike-related activities, (d-g) burst-related activities and (h-i) network burst features from DIV 4-42. Data are presented as group means ± 95% confidence interval. \**p* < 0.05.

### Characterization of burst and network burst profiles in isogenic CLN3 networks

In addition to early increases in spike rate (DIV 4-10), CLN3 neurons had higher burst rates than CLN3-Cor neurons (mean CLN3: 4.45 burst/min; CLN3-Cor: 2.52 burst/min) during the early days of maturation (DIV 4-12; Fig. 4d). Burst rate peaked at 7.53 burst/min (DIV 11) and 7.87 burst/min (DIV 39) in CLN3 and CLN3-Cor neurons respectively. As the culture matured further, CLN3-Cor neurons were bursting more frequently than CLN3 neurons during DIV 14-20 (mean CLN3: 3.89 burst/min; CLN3-Cor: 6.21 burst/min) and DIV 27-42 (mean CLN3: 2.9 burst/min; CLN3-Cor: 6.29 burst/min; Fig. 4d). Despite changes in burst rate, burst duration was similar between CLN3 and CLN3-Cor neurons during the early days of maturation (DIV 4-10; mean CLN3: 0.12 s; CLN3-Cor: 0.08 s), however, from DIV 11 onward, CLN3 neurons had shorter bursts than CLN3-Cor neurons (mean CLN3: 0.27 s; CLN3-Cor: 1.12 s; Fig. 4e).

Additionally, the percentage of spikes transformed into bursts were considerably lower in CLN3 neurons than in the CLN3-Cor neurons at most cultured time points, indicating that the spiking activity of CLN3-Cor neurons led to bursts more often than in CLN3 neurons (Fig. 4f). During DIV 7-9 CLN3 neurons had a higher percentage of spikes in bursts (mean: 36.06%) than in CLN3-Cor neurons (mean: 27.20%), however, from DIV 12 onward, CLN3-Cor neurons (mean: 75.98%) had a higher percentage of spikes in bursts than CLN3 neurons (mean: 33.38%) indicating increased activity in CLN3-Cor neurons (Fig. 4f). The percentage of spikes in bursts peaked at 50.26% (DIV 25) and 88.69% (DIV 19) in CLN3 and CLN3-Cor neurons respectively followed by a general decline in both lines toward DIV 42 (CLN3: 6.29%; CLN3-Cor: 41.21 %; Fig. 4f).

The time between bursts, known as the interburst interval (IBI), of CLN3 neurons peaked at DIV 18 (20.45 s) before decreasing toward DIV 42 (6.28 s; Fig. 4g). In CLN3-Cor neurons, the IBI surged briefly from DIV 4 (7.56 s) to DIV 8 (24.14 s), followed by a general declining trend toward DIV 42 (9.19 s; Fig. 4g). In association with an increased burst rate, the IBI was shorter in CLN3 neurons than in CLN3-Cor neurons (mean CLN3:14.61 s; CLN3-Cor: 21.96 s) during DIV 7-10 (Fig. 4g). As the culture matured during DIV 14-25, longer IBI (mean CLN3: 17.85 s; CLN3-Cor: 11.73 s) occurred concurrently with the lower burst rate in CLN3 neurons compared to CLN3-Cor neurons (Fig. 4g).

In CLN3 neurons, the network burst rate, defined as the rate of synchronous network activity between ≥ 3 electrodes per well per minute, peaked at DIV 24 (1.24 burst/min) and subsequently showed a general declining trend toward DIV 42 (0.10 burst/min; Fig. 4h). CLN3-Cor neurons generated network bursts more frequently than CLN3 neurons from DIV 9 onward (mean CLN3: 0.48 burst/min; CLN3-Cor: 2.00 burst/min), where the CLN3-Cor neuronal network burst rate peaked at DIV 15 (3.09 burst/min) before declining toward DIV 42 (1.79 burst/min; Fig. 4h). Although CLN3 and CLN3-Cor neurons demonstrated an increasing trend in network burst duration during DIV 4-34 (peak: 2.42 s (CLN3 DIV 33); 6.52 s (CLN3-Cor DIV 34)), network burst duration was longer in CLN3-Cor neurons than in CLN3 neurons from DIV 9 to DIV 42 (mean: CLN3: 1.04 s; CLN3-Cor: 3.48 s; Fig 4i). The higher burst and network burst activity in CLN3-Cor neurons for most of the culture period indicate that these cultures are more mature in neuronal development.

### Alterations in lysosomal, axon guidance and endocytosis pathways in CLN3 neurons

CLN3 is associated with various cellular processes, and therefore CLN3 deficiency is likely to have broad effects. We explored global protein changes due to CLN3 mutation using proteomics analysis. Quantitative proteomic analysis of protein extracts from isogenic CLN3 neurons was conducted at time points corresponding to early and late maturation of neurons. Differential expression analysis identified 2315 differentially expressed proteins at DIV 14 (1567 up-regulated proteins and 748 down-regulated proteins) and 1785 differentially expressed proteins at DIV 42 (1118 up-regulated and 667 down-regulated proteins). We investigated the effect of independent neuronal batches, genotype and DIV on the proteome using PCA (Fig. 5a). The PCA plot revealed clustering of biological replicates of CLN3 and CLN3-Cor neurons, but separation according to genotype and DIV. DIV explained 60% of the variation between samples (PC1) while genotype explained 26% of the variation (PC2). Bioinformatic analysis was then used to identify KEGG pathways that were enriched in the sets of differentially abundant proteins. Significantly enriched pathways represented by the proteins detected at reduced levels in CLN3 neurons (DIV 14 and DIV 42) included endocytosis and axon guidance, while the lysosomal pathway was over-represented among the proteins detected at higher levels in CLN3 neurons (Fig. 5b). The subsets of proteins related to lysosome, axon guidance and endocytosis pathways are represented in Fig. 6a, b, c according to their abundance ratios (CLN3/CLN3-Cor).

**Fig. 5.**
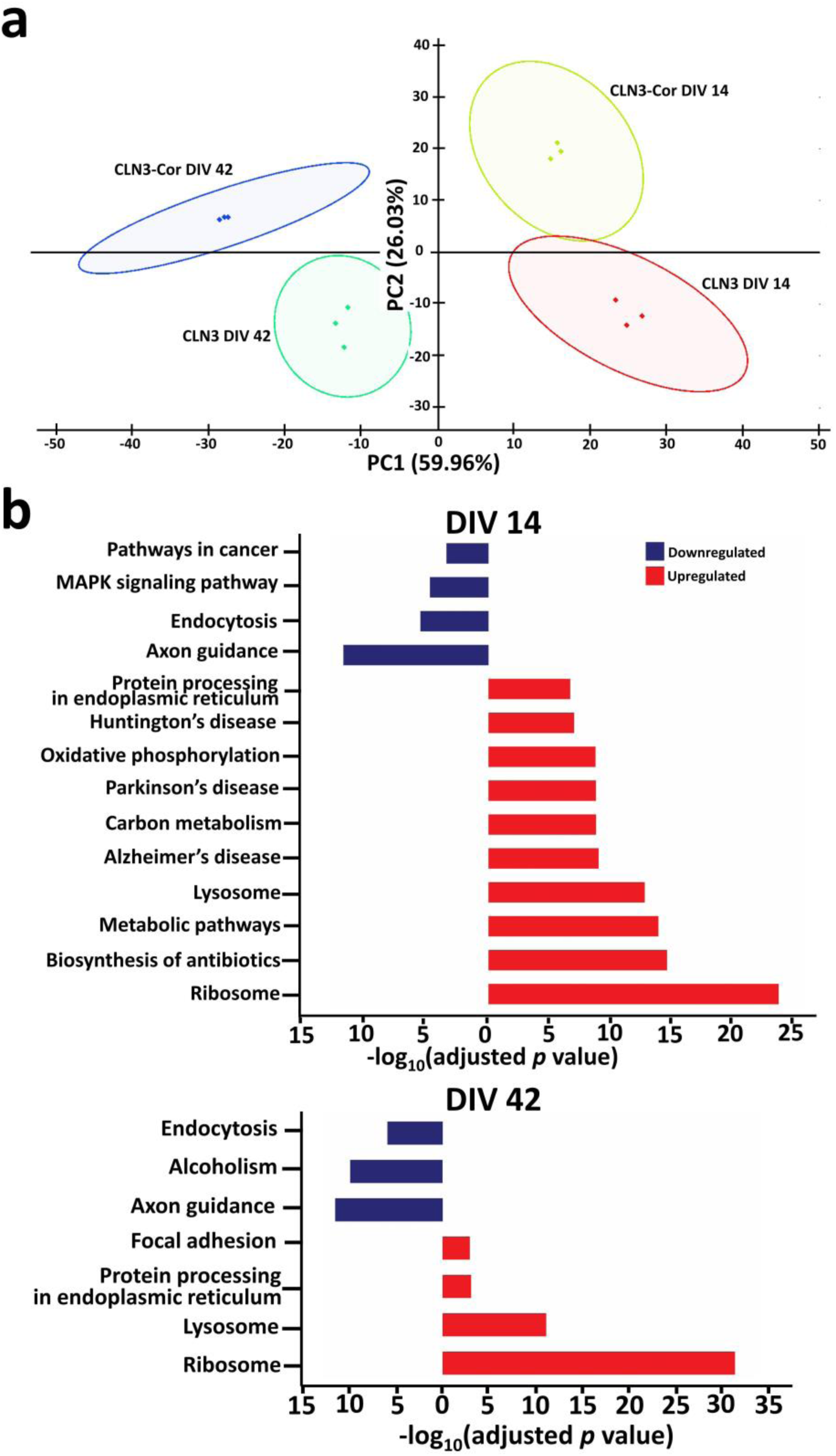
Proteomic profile of isogenic CLN3 neurons. (a) PCA plot of CLN3 and CLN3-Cor neurons at DIV 14 and 42. b) Enriched KEGG pathways in CLN3 neurons at DIV 14 and DIV 42 filtered based on thresholds (protein count ⩾ 30 and -log10(Benjamini adjusted *p* value ≥1.3)).

**Fig. 6.**
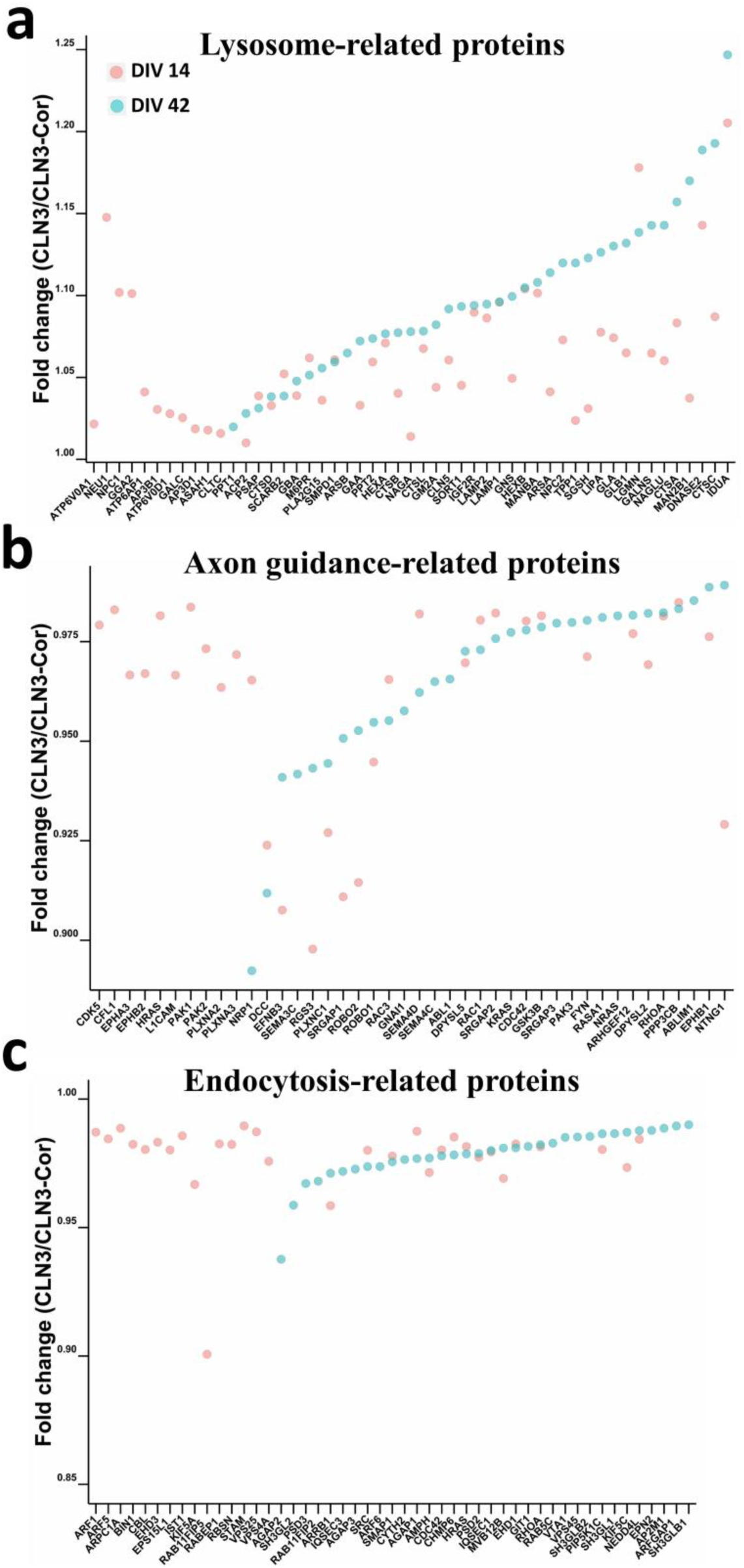
Altered KEGG pathways in CLN3 neurons. Overview of mean fold change of protein expression in CLN3 neurons compared to CLN3-Cor neurons for enriched KEGG pathways related to (a) lysosomal proteins increased in abundance (mean fold change >1) (b) axon guidance-related proteins reduced in abundance (mean fold change <1) and (c) endocytosis-related proteins reduced in abundance (mean fold change <1) at DIV 14 and 42 (n=3 independent cultures per time point).

### LAMP1 hyperglycosylation, altered ultrastructure of storage material and endocytosis in CLN3 neurons

To further explore lysosomal alterations in CLN3 neurons, we performed Western blotting for detection of the lysosomal marker protein LAMP1. CLN3-Cor and genetically unrelated healthy control iPSC-derived neurons showed a single diffuse protein band of ∼90 kDa. In contrast, an additional band of ∼120 kDa was identified in CLN3 neurons (Fig. 7a). The total level of LAMP1 was higher in CLN3 neurons than in CLN3-Cor neurons at DIV 28 and 42 (Fig. 7b). The ratio of LAMP1 120 kDa/90 kDa bands showed a decreasing trend from DIV 14 to DIV 42 (Fig. 7c). To identify the type of posttranslational modification that may be related to the larger LAMP1 protein, protein extracts from CLN3 and CLN3-Cor neurons were treated with N-glycanase and endoglycosidase H. Western blot analysis showed a smaller band of ∼40 kDa in size after N-glycanase digestion (Fig. 7d). Meanwhile, Western blot analysis of endoglycosidase H treatment showed a diffuse band of the larger-sized LAMP1 band in CLN3 neurons with full digestion of the 90 kDa LAMP1 form (Fig. 7d). This shows that the posttranslational modification of the larger LAMP1 protein involves complex or hybrid N-linked glycosylation.

**Fig. 7.**
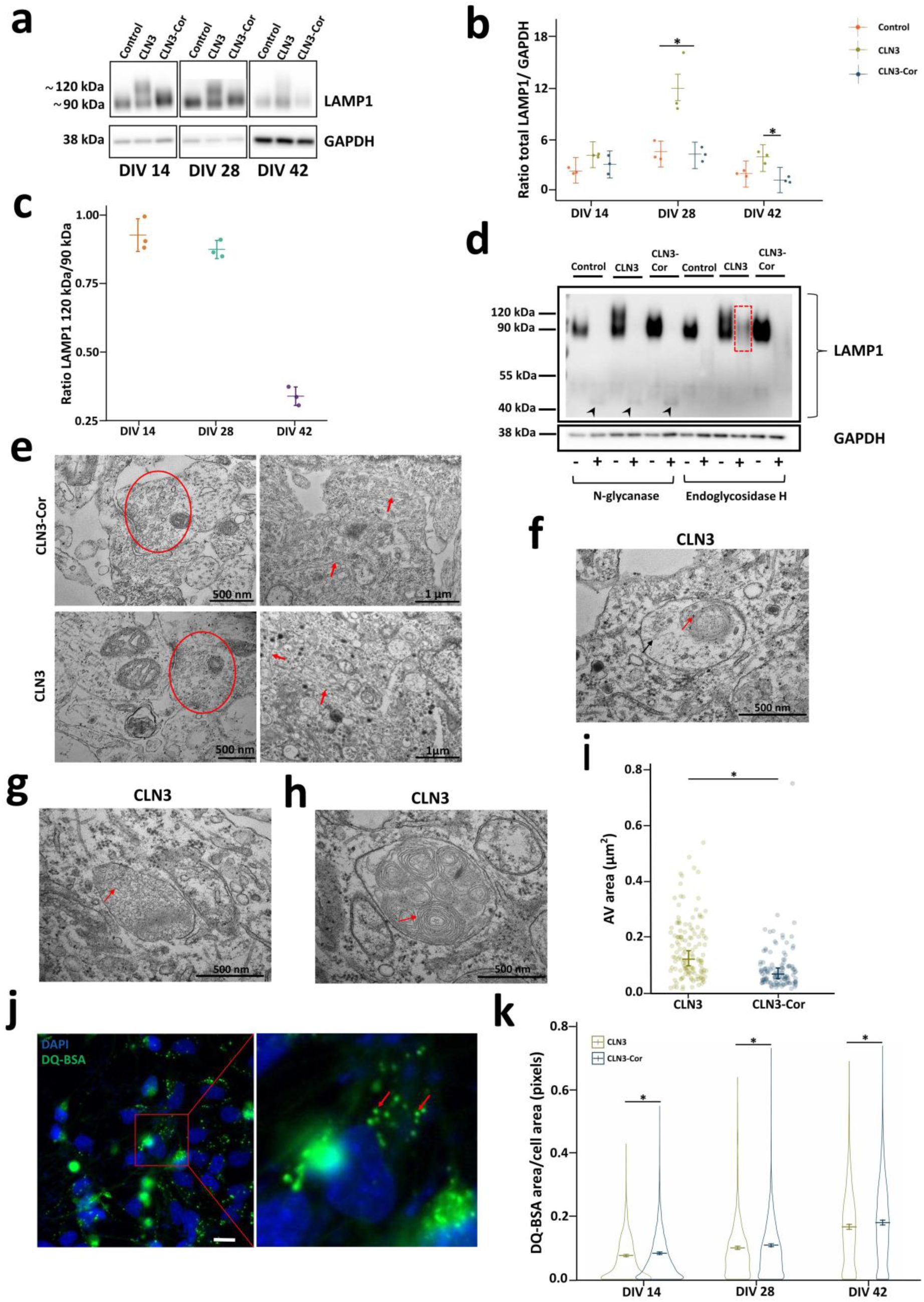
Alterations in LAMP1 expression, storage material and endocytosis. (a) Representative western blots showing LAMP1 protein expression across different time points, DIV 14, 28 and 42. Bands corresponding to the higher molecular weight (∼120 kDa) form of LAMP1 protein were present only in the CLN3 neurons. (b) Expression of total LAMP1 in all cell lines. (c) LAMP1 upper band/lower band expression level ratio in CLN3 neurons across different time points. Data are represented as mean and standard deviation. (d) LAMP1 protein expression after treatment with or without N-glycanase and endoglycosidase H digestion. The N-glycanase-deglycosylated form of LAMP1 is highlighted by arrowhead. Diffuse LAMP1 band in CLN3 neurons after endoglycosidase H digestion is highlighted by red dashed outline. (e) Representative images demonstrating neuronal phenotype of both cell lines, including synaptic vesicles (red outline) and neurofilaments (red arrows). Different storage materials were observed in the CLN3 culture, including (f) osmiophilic (red arrow) and fibrillar deposits (black arrow), (g) curvilinear structures (red arrow) and (h) multilamellar structures (red arrow). (i) Quantitation of autophagic vacuole (AV) area at DIV 42 (j) Representative fluorescence image showing DQ-BSA appearing as bright green puncta (indicated by inset) in the cytoplasm in CLN3-Cor neurons (left panel), further zoomed-in and indicated by red arrows (right panel). (k) DQ-BSA area in isogenic CLN3 neurons across different time points. All data, unless otherwise specified, are presented as group means ± 95% confidence interval (n=3 independent cultures per time point). \**p* < 0.05.

As CLN3 mutation is associated with accumulation of subunit c of mitochondrial ATP synthase in cells, the subunit c levels in neuronal culture were examined through Western blot. Quantitation of subunit c protein levels in the isogenic CLN3 neurons did not show differences through DIV 14, 28 and 42 (Additional file 8: Figure S7). Additionally, we further evaluated the ultrastructure of storage material in the isogenic CLN3 neurons through electron microscopy. Neurons from both genotypes have comparable neuronal phenotypes in terms of formation of synaptic vesicles and neurofilaments (Fig. 7e). There were numerous autophagic vacuoles with a size range of 500 - 1000 nm that were filled with heterogeneous storage material including osmiophilic and floccular deposits (Fig. 7f), curvilinear structures (Fig. 7g) and multilamellar structures (Fig. 7h) in CLN3 cells. The majority of storage material observed was in the form of multilamellar structures. However, fingerprint deposits were not observed in CLN3 cells. Quantitation of the area of autophagic vacuoles revealed an increased autophagic vacuole area in CLN3 cells compared to CLN3-Cor cells (Fig. 7i).

Previous studies have demonstrated impairment of endocytosis in CLN3-deficient models [19, 38]. To investigate the effect of CLN3 mutation on endocytosis, we performed a DQ-BSA assay, where formation of fluorescent DQ-BSA puncta indicates active endocytic trafficking process (Fig. 7j). Statistical tests comparing the DQ-BSA area in both cell lines revealed a minor but significant decrease in DQ-BSA area in CLN3 neurons across all time points (Fig. 7k).

## DISCUSSION

The patient-specific isogenic CLN3 neuronal cell model in this study was found to recapitulate features of CLN3 mutation, including upregulation of lysosomal protein expression and increased intracellular storage materials. In addition, this study provides new insight into LAMP1 hyperglycosylation and the electrophysiological alterations in CLN3 neurons from day 4 to 42 of neuronal maturation using MEA.

Using MEA, we analyzed selected features of spikes, bursts, and network events to compare the functional activity over time from day 4 to day 42 after the initiation of neuronal maturation in isogenic CLN3 neurons. While spikes and bursts reflect the overall activity of the neuronal network, network burst events demonstrate the synchronicity in spikes and bursts within the neuronal network [42]. The MEA findings showed that CLN3 and CLN3-Cor neuronal networks demonstrated similar phases of development in neuronal activity patterns. CLN3-Cor and CLN3 neuronal networks fired spikes and bursts and upon maturation, increased connectivity and synchronicity was observed within network bursts, however, the time scale of development and levels of activity varied between cell lines. Although the CLN3 neurons increased their activity over the first 10 days of maturation, their activity either plateaued or increased gradually toward the end of the culture period. This is in contrast to CLN3-Cor neuronal activity, which showed a consistent increase in activity from early days of maturation to later time points of the culture period [45].

Network bursts play an important role in nervous system development and maturation, and are strongly correlated with the strengthening of synapses [26, 69], both glutamatergic and GABAergic [27]. The CLN3 neurons showed considerably lower synchronicity within the network compared to CLN3-Cor neurons, as indicated by lower network burst rate and duration in CLN3 neurons. Additionally, during the later stage of maturation (DIV 28), CLN3 neurons had lower expression of several genes related to glutamatergic and GABAergic neurons and synaptic protein, potentially indicating altered maturation during development of the neuronal culture. This reduced development and maturation of neuronal networks in CLN3 neurons could be related to neurodevelopmental aspects of CLN3 disease, where transcription factors involved in central nervous system development have been reported to be downregulated in CLN3-derived organoids [23]. Our proteomics data supported the neurodevelopmental alterations in CLN3 neurons, showing the downregulation of axon guidance-related proteins in CLN3 neurons. The downregulated proteins in axon guidance comprised several important classes of axon guidance molecules and their receptors. These include NTNG1 guidance cue which binds to the transcription activator DCC [58], L1CAM (cell adhesion molecule) which is essential for axonal elongation and dendritic arborization [53], ROBO1 and ROBO2 receptors which guide major forebrain axonal [37] projections and SEMA4D, a semaphorin which repels axons [57]. Formation of the central nervous system during embryogenesis requires neurons to extend axons to distant targets for synaptic transmission and to develop functional circuits [52]. Disruption to axon guidance could therefore affect neuronal differentiation and maturation, potentially explaining the lower levels of firing and network burst activity in the CLN3 neurons. Difficulty in differentiation of CLN3 neural progenitor cells (homozygous CLN3 966 bp deletion) using the monolayer method has been reported, which was then remedied by formation of neuralized embryoid bodies [36]. Severe developmental failure in CLN3^Q352X^ organoids upon initiation of differentiation has also been described, however, in the CLN3^Q352X^ organoids which did differentiate normally, the expression of neuronal markers was similar to control organoids [23]. These and our findings suggest that the mutation of CLN3, which has a functional role in neurogenesis during early embryonic development [41], may affect neuronal differentiation and maturation to a variable extent depending on the type of pathogenic variant.

In addition to axon-related alterations, our proteomics analysis also identified altered endocytosis-related proteins at both DIV 14 and 42. Impaired endocytosis has been reported in CLN3 patient fibroblasts, CLN3-deficient yeast [9], CLN3 mouse cerebellar cells [19] and CLN3 endothelial cells [66]. In the CLN3 neurons, proteins related to clathrin-mediated endocytosis, which is essential for synaptic plasticity, neurotransmission [31] and axonal development for pathfinding [73] were downregulated. Data from the DQ-BSA Green experiment indicate that CLN3 mutation may have a minor effect on endocytosis, possibly due to residual function of CLN3 E295K protein which compensates for the deficiency of CLN3 protein from the 966 bp deletion allele. The E295K and other missense variants are trafficked to lysosomes, whereas the 966 bp deletion and other frameshifting and nonsense mutations cause CLN3 to be retained within the endoplasmic reticulum [24].

In this study, upregulation of lysosome-related proteins at DIV 14 (49 proteins) and DIV 42 (40 proteins) was observed in accordance with previous research, which demonstrated significant elevation of lysosomal proteins SCARB2, HEXB and TPP1 in CLN3 mouse brain [63], HEXA and LAMP1 in urine of CLN3 patients [28], PPT1 and GLA in lysosomal fractions of CLN3^*Δex7/8*^ mouse cerebellar cells [59], and of PPT1, TPP1, NAGA and NAGLU in CLN3 brains following autopsy [62]. The changes in lysosomal protein abundance may be a compensatory cellular response to the accumulation of various substrates within the lysosome. Amongst the lysosomal proteins which were upregulated with higher fold change at DIV 42 compared to DIV 14 were CTSC, TPP1, NAGA and GM2A. The elevation of these proteins in our study indicates possible increase in the catalysis of substrates such as gangliosides by GM2A, HEXA and HEXB [12], protein by TPP1 and CTSC, and glycoprotein by NAGA to prevent the accumulation of these substrates. Notably, the accumulation of GM2A within lysosomes of neurons which occurs in Tay-Sachs disease, another lysosomal storage disorder, has led to neuronal death [16]. A separate study in *Cln3*^*Δex7/8*^ mice has also reported accumulation of GM3 gangliosides due to altered expression of ganglioside metabolizing enzymes [64]. TPP1 has been suggested to be involved in the degradation of subunit c of mitochondrial ATP synthase [17]. Therefore, elevation of TPP1 activity, which has been described in CLN3 mouse brain [61] and CLN3 patient brains [29, 61] may be important to prevent accumulation of subunit c. Additionally, alterations to lysosomal proteins may also have an impact on axon growth, where inhibition of lysosome transport to the distal axon alters the size and dynamics of growth cone, which could be related to disruption of lysosome-mediated delivery of signaling and adhesion molecules [18] or local degradation of cargos in axons to maintain axon homeostasis [39]. Together, these results suggest that CLN3 deficiency alters lysosomal function and endocytosis, and that this may have an impact on the synaptic transmission and axonal growth, as indicated in our MEA and proteomics studies.

The accumulation of osmiophilic deposits, suggestive of lipid storage in our CLN3 neuronal model, was also observed in *CLN1* (PPT1) mutation. CLN1 deficiency and other lipid storage disorders such as Gaucher disease are associated with the accumulation of saposin A and D, which degrade sphingolipids [67]. Meanwhile, the curvilinear storage materials observed in CLN3 neurons are the dominant ultrastructure of storage bodies in CLN2 mutation. The curvilinear storage materials comprise mainly proteins and are associated with subunit c of mitochondrial ATP synthase, a highly hydrophobic and major accumulating protein in CLN2 mutation [21, 48]. The accumulation of multilamellar membranous whorls in the storage contents of CLN3 neurons are also suggestive of disruption in lipid degradation [50]. The presentation of heterogeneous storage materials in CLN3 neurons highlights the possibility of deficit in degradation of multiple types of cellular wastes comprising protein, lipid and glycan. The appearance of heterogeneous storage materials observed in CLN3 neurons, rather than the predominant fingerprint profile observed in typical CLN3 mutations suggest that this is a CLN3 variant or cell line specific ultrastructural phenotype which will require confirmation from other CLN3 models. In line with data from our proteomics study, Western blot analysis of total LAMP1 protein level revealed higher expression of LAMP1 protein in CLN3 neurons than in CLN3-Cor neurons at DIV 28 and 42. Together, these data further support our findings of lysosomal alterations in CLN3 neurons, evidenced by upregulation of lysosomal proteins in the global proteome and increased accumulation of storage materials at the electron microscopy level.

The hyperglycosylated LAMP1 observed in CLN3 neurons was not apparent in both CLN3-Cor and genetically unrelated healthy control iPSC-derived neurons. The higher molecular weight form of LAMP1 was sensitive to N-glycanase digestion but resistant to endoglycosidase H deglycosylation, indicating the presence of complex N-linked glycans. To our knowledge, our study is the first to report the occurrence of LAMP1 hyperglycosylation in CLN3 mutation. A similar finding has been observed in *Npc*^-/-^ mice, a model of Niemann-Pick disease (NPC) type C, and which has been associated with Purkinje neuron loss [8]. The hyperglycosylation of LAMP1 in our CLN3 neuronal model occurred concurrently with altered levels of NPC1 and NPC2 as observed in our proteomics study. NPC1 and NPC2 are lysosomal glycoproteins which are involved in egress of cholesterol from lysosomes to the cytoplasm [7, 43]. Hyper-glycosylated LAMP1 has been shown to impair transportation of cholesterol from the lysosome to the endoplasmic reticulum in NPC1 protein-deficient cells [35]. Therefore, LAMP1 hyperglycosylation and increased levels of NPC1 and NPC2 in the CLN3 neurons could be associated with accumulation of cholesterol in the lysosomes [35]. This distinct hyperglycosylated form of LAMP1 was observed predominantly in CLN3 neurons at DIV 14, 28 and to a lesser extent in the later time point at DIV 42. The reduced hyperglycosylation toward DIV 42 could be a compensatory response to increase efflux of accumulated contents in the lysosome, for example, cholesterol. Substrate accumulation within the lysosome, especially subunit c of mitochondrial ATP synthase, is a hallmark feature of CLN3 disease [48]. However, Western blotting of subunit c in isogenic CLN3 neurons did not reveal altered expression levels of subunit c in CLN3 neurons suggesting that subunit c accumulation may be occurring further downstream in the pathological process.

## CONCLUSION

Findings from this study further our understanding of cellular alterations in CLN3 disease, especially the lysosomal pathway and neurodevelopmental aspects, caused by compound heterozygous mutations associated with an atypical, protracted phenotype. Our study of early onset and subtle phenotypes in isogenic neurons could potentially aid in drug discovery to halt disease progression before neurodegeneration occurs, and which may be applicable to other CLN3 genotypes in addition to that studied here. In particular, modulation of key proteins in the lysosomal pathway may be a potential target to protect against neurodegeneration. For example, trehalose, a transcription factor EB activator which promotes lysosomal clearance of proteolipid aggregates [49], in combination with miglustat is being investigated in a clinical trial for its efficacy in the treatment of CLN3 disease (NCT05174039). Future studies using iPSC-derived neuronal models that capture a diversity of CLN3 genotypes will help to uncover these drug targets.

## DECLARATIONS

### Ethics approval

The use of human biological material for research purpose of this study was approved by the Human Research Ethics Committee of Tasmania (H0014124). Work involving genetic manipulation was approved by the University of Tasmania Institutional Biosafety Committee.

## Acknowledgements

We are grateful to our participant and their family for providing a skin biopsy. We would like to thank the Batten Disease Support and Research Association Australia (RGP204), and the Royal Hobart Hospital Research Foundation for funding this project. We thank Olivier Bibari (University of Tasmania) and Helena Liang (Centre for Eye Research Australia) for technical assistance.

## Author’s contributions

SC, SP, JT, AK, AH and AC contributed to the conception and design of the study; SC and RW performed the experiments, SC, RW and AB contributed to data analysis; SC, RW and SP wrote the manuscript. TW, AG, JCV, AP, and JBR provided resources. All the authors revised the manuscript and agreed to the final version of the manuscript.

