## Supplemental Files for "Lysosomal alterations and decreased electrophysiological activity in CLN3 disease (966 bp deletion, E295K) patient-derived cortical neurons"

**ADDITIONAL FILES**

1. **Additional file 1**

**Table S1: List of all oligonucleotides used in this study**

| **Oligo name** | **Sequence** | **Region to amplify** | **Expected PCR band size** |
|---|---|---|---|
| EBNA-1 F | ATCGTCAAAGCTGCACACAG | Sequence in EBNA-1 episomal vector | 666 bp |
| EBNA-1 R | CCCAGGAGTCCCAGTAGTCA | | |
| CLN3 F1 | GGATGAATTAGATGGAGATTGAGG | CLN3 DNA: region spanning exons 7,8 | 966 bp deletion: 563 bp<br><br>Wild type: 1529 bp HDR insertion with puromycin cassette: 2581 bp<br><br>HDR insertion without puromycin cassette: 1563 bp |
| CLN3 R1 | CTCATCCTACTTCTAATCACCTTG | | |
| CLN3_SS_cdna_F1 | TCTGTCTCTACGGCTGCTGTGC | CLN3 cDNA: region spanning exons 7,8 | Deleted: 578bp<br><br>Undeleted: 795bp |
| CLN3_SS_cdna_R1 | GAACACCAGGTTGAGGCACTGC | | |

**Table S2 List of Taqman assay probes for neural induction and differentiation**

| Marker | Gene name | Taqman Assay ID |
|---|---|---|
| Pluripotency | *OCT4* | Hs01895061_u1 |
| Pluripotency | *NANOG* | Hs00415716_m1 |
| Neuroepithelial stem cell | *NESTIN* | Hs04187831_g1 |
| Neuroepithelial stem cell | *PAX6* | Hs01088114_m1 |
| Mature neuron | *MAP2* | Hs00258900_m1 |
| Microtubule | *TUBB3* | Hs00801390_s1 |
| γ-aminobutyric acid biosynthesis enzyme | *GAD1* | Hs01065893-m1 |
| γ-aminobutyric acid biosynthesis enzyme | *GAD2* | Hs00609534_m1 |
| α-amino-3-hydroxy-5-methyl-4-isoxazolepropionic acid (AMPA) receptor | *GRIA2* | Hs00181331_m1 |
| N-methyl-D-aspartate (NMDA) receptor | *GRIN1* | Hs00609557_m1 |
| Excitatory amino acid transporter | *SLC1A7* | Hs00220404_m1 |
| Excitatory amino acid transporter | *SLC1A2* | Hs01102423_m1 |
| Excitatory amino acid transporter | *SLC1A3* | Hs00188193_m1 |
| Postsynaptic density protein | *DLG4* | Hs01555370_g1 |
| Ceroid-lipofuscinosis, Neuronal 3 | *CLN3* | Hs00164002_m1 |
| Reference gene | *EEF2* | Hs00157330_m1 |

**Table S3: MEA parameters and description**

| **Parameters** | **Description** |
|---|---|
| Spike rate (Hz) | Number of spikes/second |
| Interspike interval (s) | Time interval between two consecutive spikes |
| Percentage of active electrode (%) | Percentage of number of active electrodes divided by total number of electrodes |
| Burst rate (burst/min) | Number of bursts per minute |
| Burst duration (s) | Duration of burst |
| Interburst interval (s) | Time interval between two consecutive bursts |
| Percentage of spikes in burst (%) | Percentage of number of spikes which were transformed into bursts divided by total number of spikes |
| Network burst rate (burst/min) | Number of network bursts per minute |
| Network burst duration (s) | Duration of network burst |

**Table S4: Off-target analysis**

| Sequence | PAM | Gene | Chromosome | Mismatches | Modification |
|---|---|---|---|---|---|
| CAGAGTATGGA CTTGAAGGA | AAG | | Chr20 | 3 | No |
| CCCGTTAGGGA CTTGAAGGA | AAG | | Chr19 | 3 | No |
| CAAGGCAGGG AATTGAAGGA | AGG | | Chr5 | 2 | No |
| GCAGGTAGAG ACTTGAAGGA | GAG | | Chr2 | 3 | No |
| CAATCTTGGGA CTTGAAGGA | AAG | | Chr12 | 3 | No |
| CTAGGCAAGGA CTTGAAGGA | GGG | | Chr15 | 3 | No |
| ACAGGTAGGG ACTTGAGGGA | AAG | ENSG00000 157429 | Chr16 | 3 | No |
| CAATGCAGGAA CTTGAAGGA | GAG | | Chr3 | 3 | No |
| CAAGGTAGGA ACTTGAAAGA | TGG | | Chr2 | 2 | No |
| CAAAGTAGGTA TTTGAAGGA | AAG | | Chr10 | 3 | No |
| CCAGGTCGGGT CTTGAAGGA | TGG | ENSG00000 279069 | Chr17 | 3 | No |
| CAAGTTTTTGA CTTGAAGGA | AAG | ENSG00000 182836 | Chr5 | 4 | No |

### 2. Additional file 2

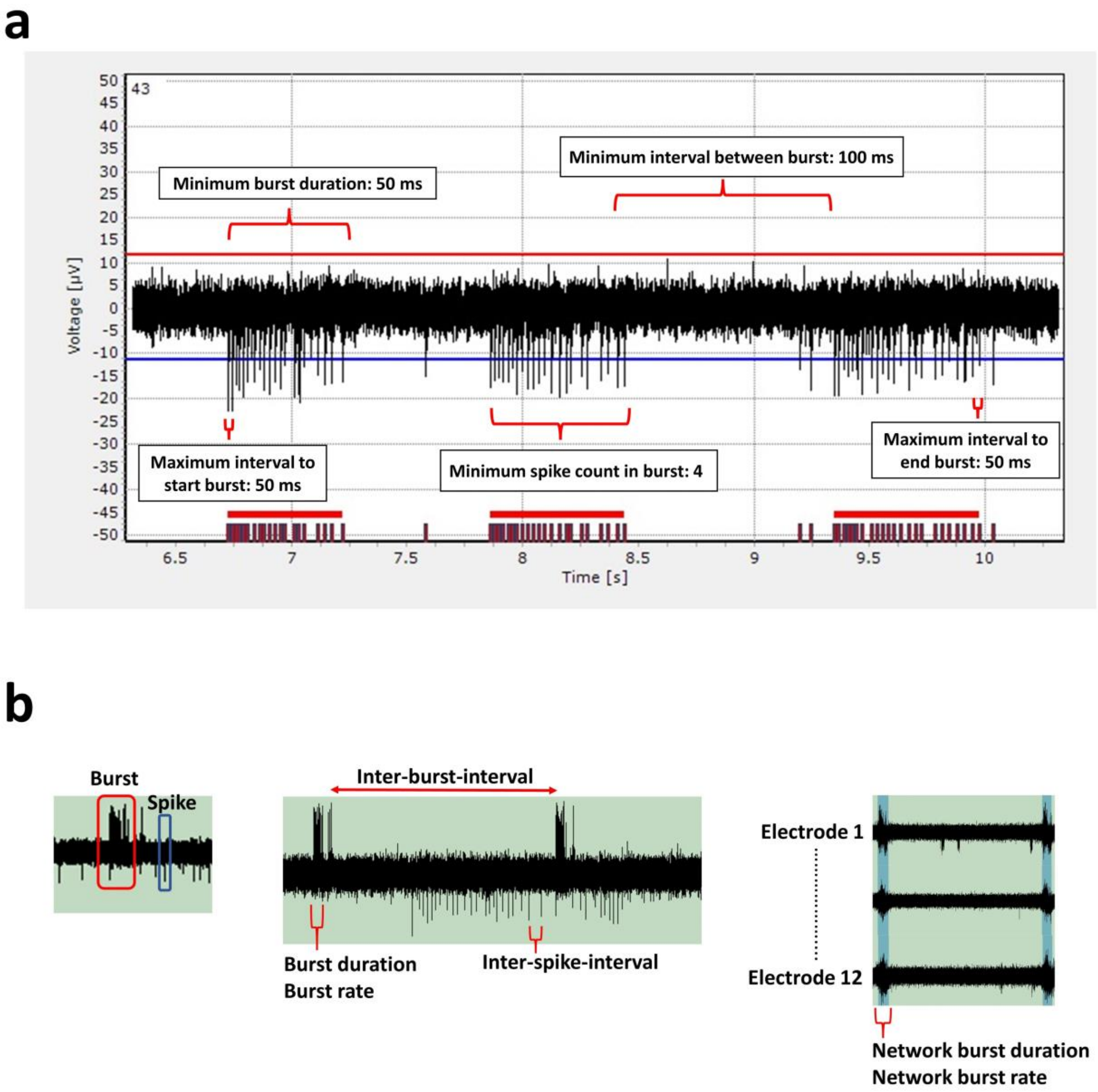


**Figure S1** MEA parameters used to measure electrophysiological activity. (a) Five threshold parameters were used to identify bursts as shown on raw data. (b) Schematic overview of extracted parameters from MEA raw data including spikes, bursts and network bursts.

#### 3. Additional file 3

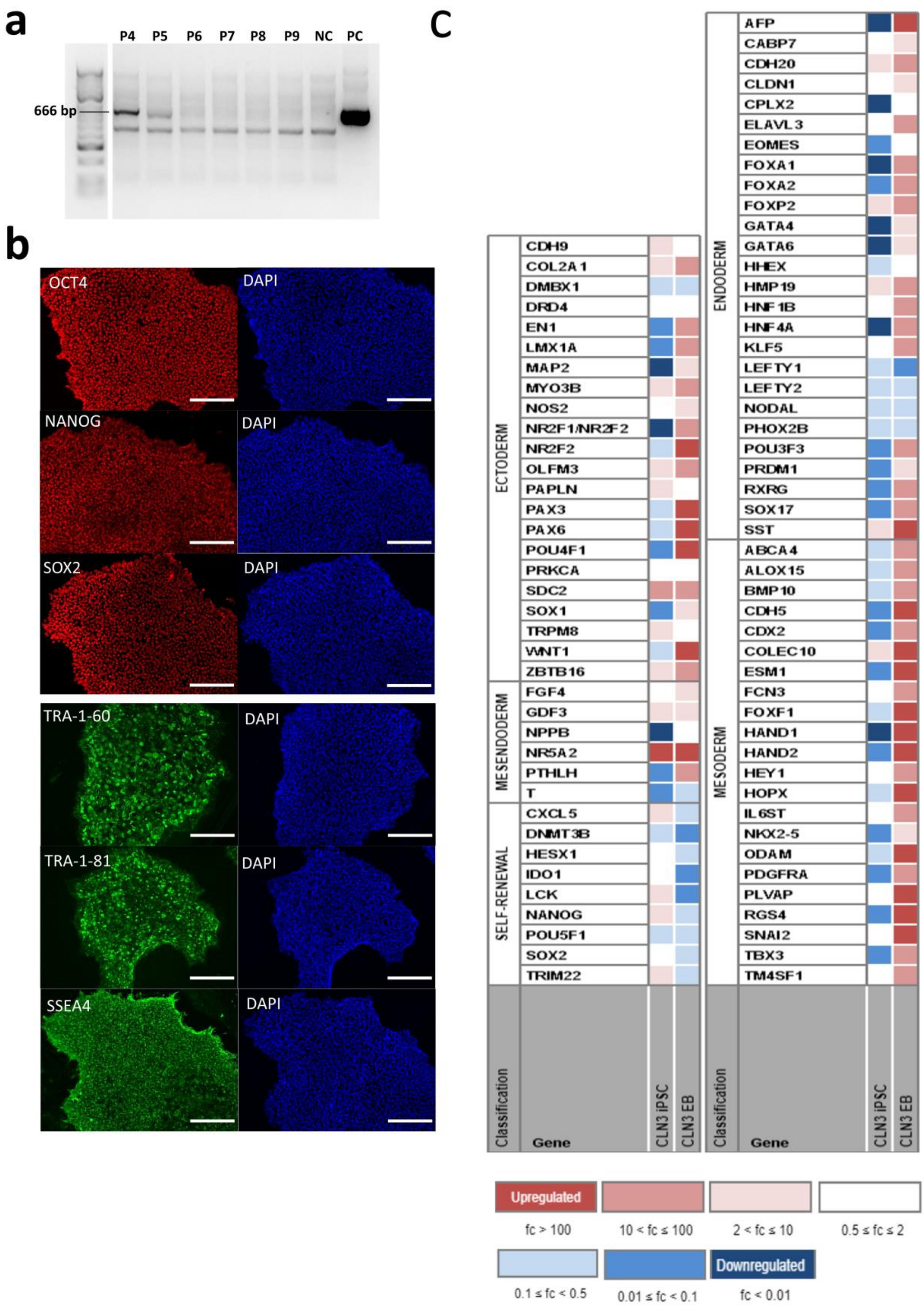


(See legend on next page.)

(See figure on previous page.)

**Figure S2** Quality control assessment of patient fibroblast-derived CLN3 iPSCs. (a) Endpoint PCR showed removal of EBNA1 vector (666 bp) from CLN3 iPSCs after several passages (P). NC and PC represent negative control from untransfected fibroblasts while PC represents positive control from EBNA1 plasmid. (b) CLN3 iPSCs displayed nuclear (OCT4, NANOG and SOX2) and surface (TRA-1-60, TRA-1-81 and SSEA-4) markers indicating pluripotency. Scale bars: 200 µm. (c) CLN3 iPSCs showed an upregulation of pluripotent genes with downregulation of germ layer genes. Meanwhile EBs demonstrated a downregulation of pluripotent genes with upregulation of germ layer genes concurrently.

**4. Additional file 4**

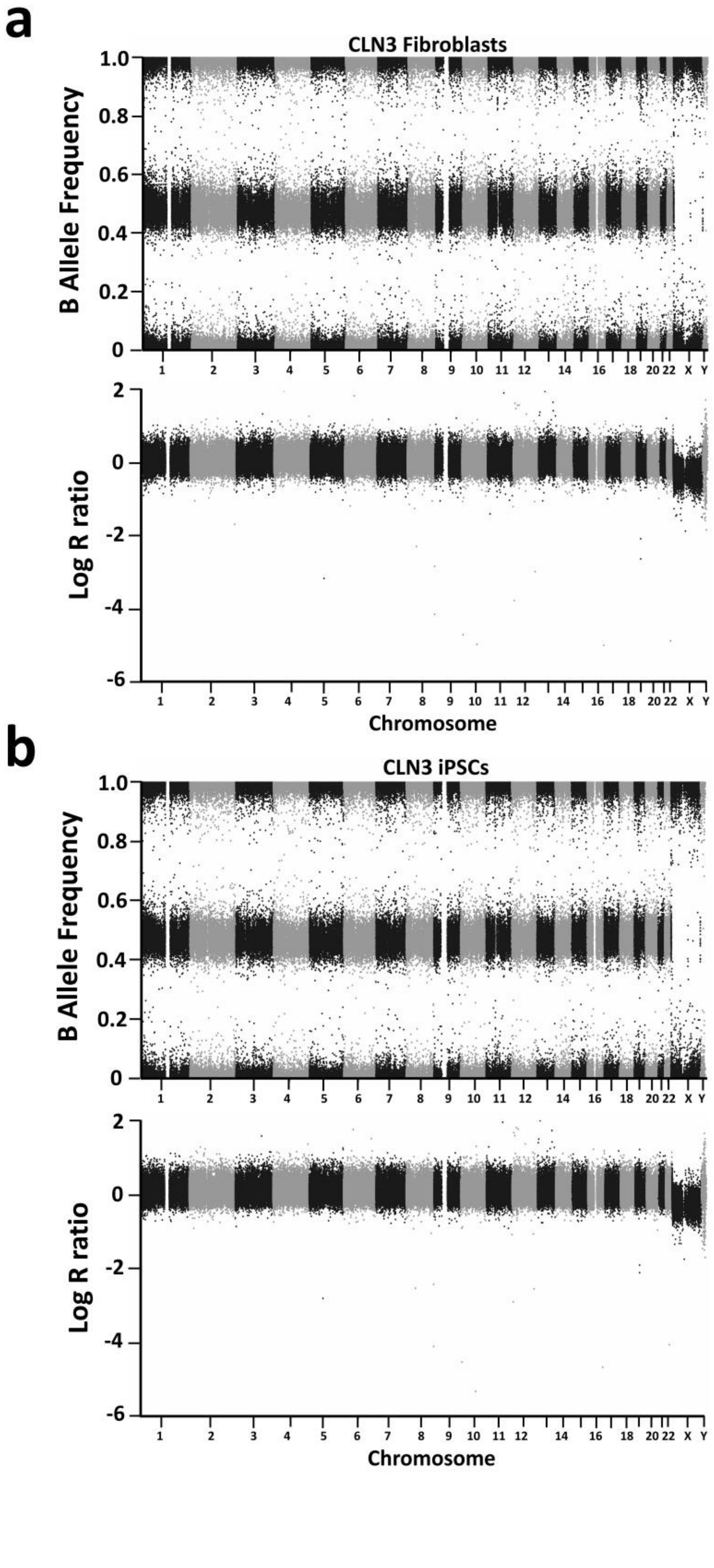


(See legend on next page.)

(See figure on previous page.)

**Figure S3 Copy number variation analysis of CLN3 iPSCs.** Top and bottom panels show B allele frequency (BAF) and Log R ratio (LRR) respectively for the (a) CLN3 parental fibroblasts (b) CLN3 iPSCs.

### 5. Additional file 5

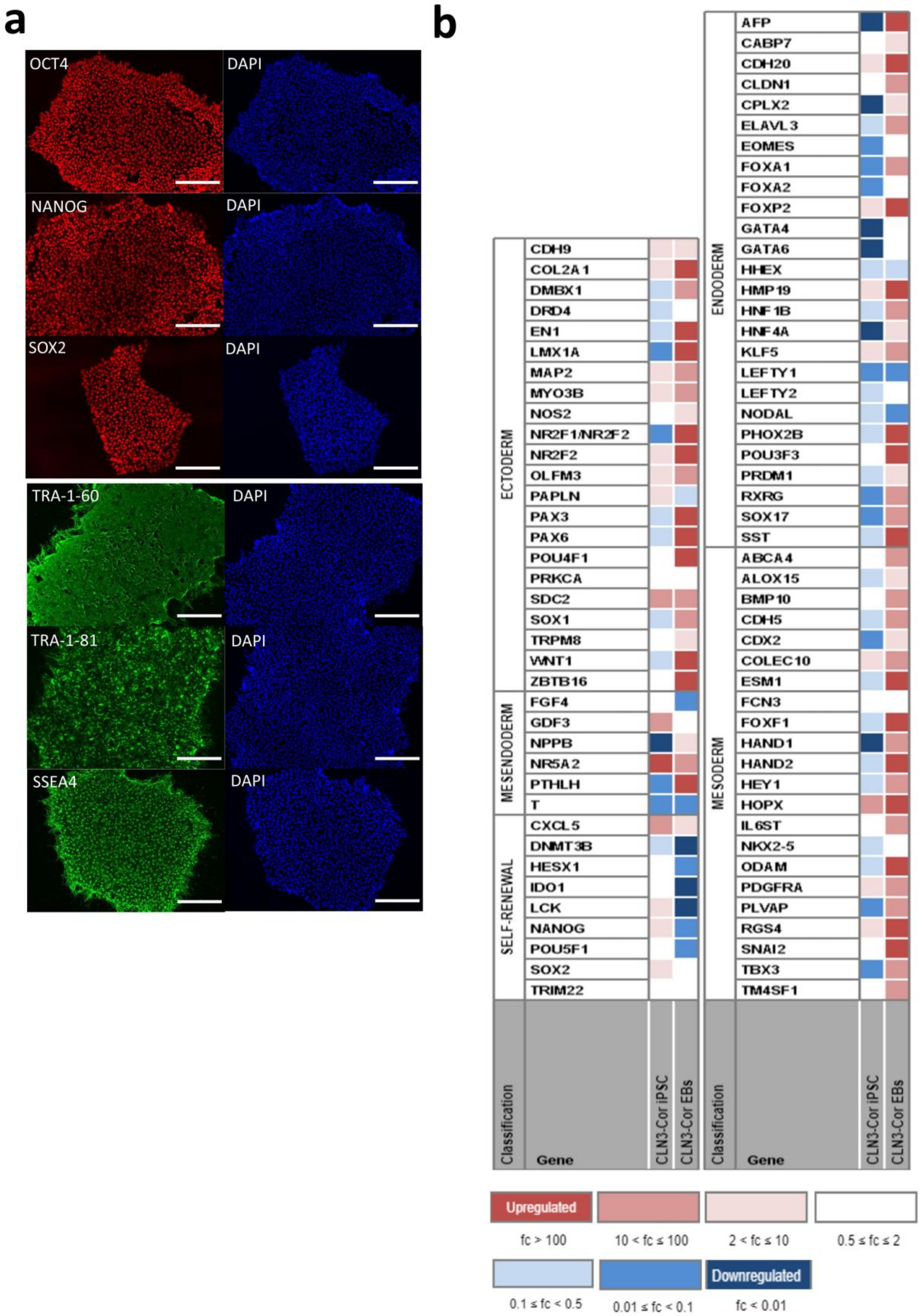


(See legend on next page.)

(See figure on previous page.)

**Figure S4** Quality control assessment of CLN3-Cor iPSCs. (a) CLN3-Cor iPSCs stained positive for OCT4, NANOG, SOX2, TRA-1-60, TRA-1-81 and SSEA4. Scale bars: 200 µm. (b) Heatmap from Taqman scorecard analysis provides an overview of the expression level of pluripotent and germ layer-specific genes in CLN3-Cor iPSCs and EBs, demonstrating CLN3-Cor iPSCs were pluripotent and the EBs were able to differentiate into 3 germ layer markers.

### 6. Additional file 6

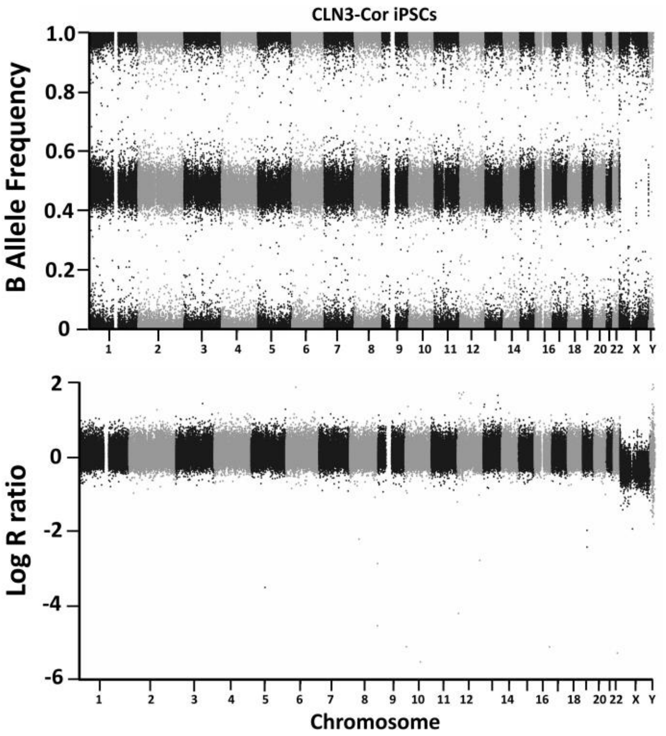


**Figure S5** Copy number variation analysis of CLN3-Cor iPSCs. Top and bottom panels show B allele frequency (BAF) and Log R ratio (LRR) respectively for the CLN3-Cor iPSCs.

### 7. Additional file 7

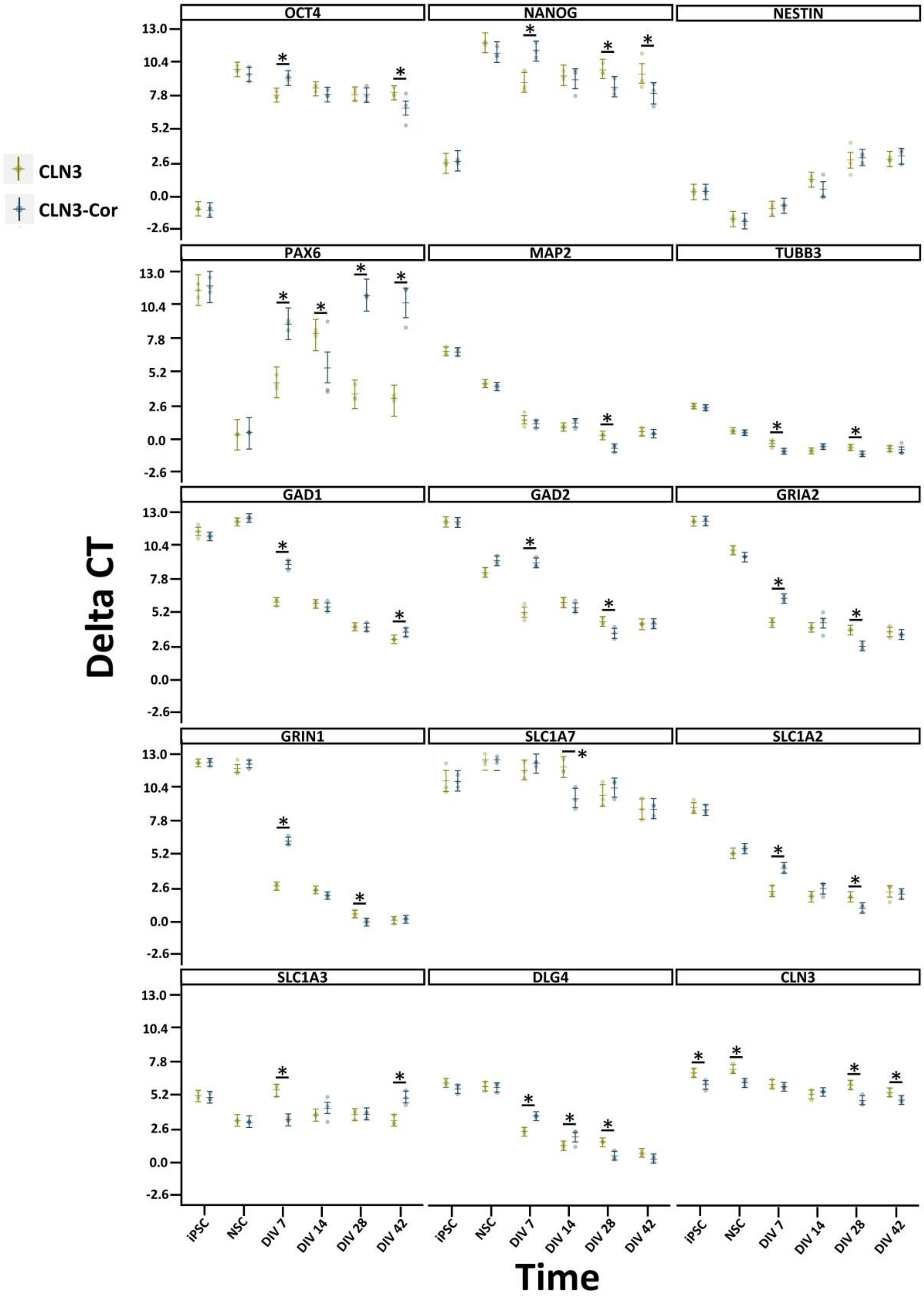


(See legend on next page.)

(See figure on previous page.)

**Figure S6** Characterization of NSCs and neurons derived from isogenic CLN3 iPSCs. mRNA expression levels of genes indicating neural induction and differentiation were examined with qPCR in iPSCs, NSC and neurons at various time points of differentiation. Data are presented as group means ± 95% confidence interval, *$p < 0.05$.

### 8. Additional file 8

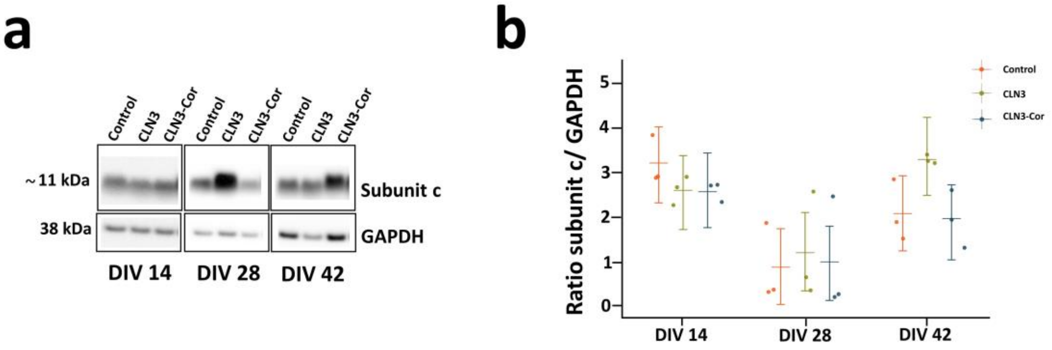


**Figure S7** Expression of subunit c in isogenic CLN3 neuronal culture. (A) Western blots showing subunit c protein expression across different time points. (B) Quantitation of subunit c in both cell lines across different timepoints. (n=3 independent cultures per time point). Data are presented as group means ± 95% confidence interval.

9. **Additional file 9: Raw images of Western Blots**

**Fig. 7a**

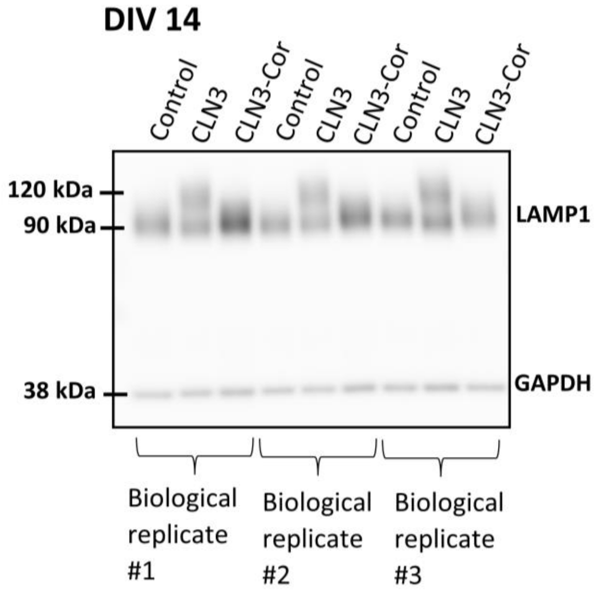


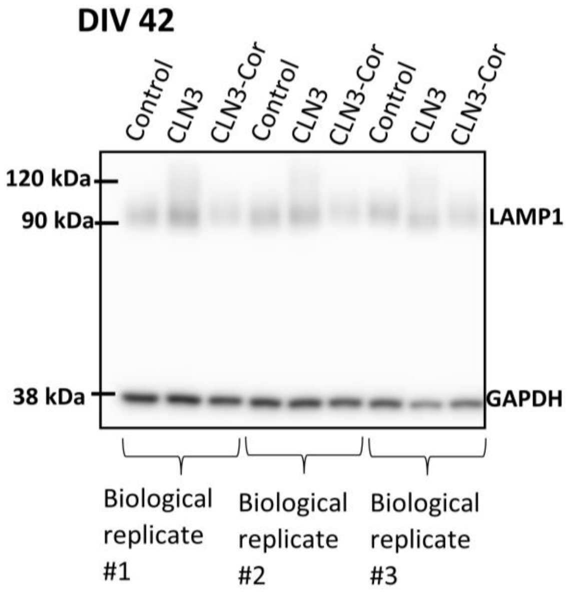


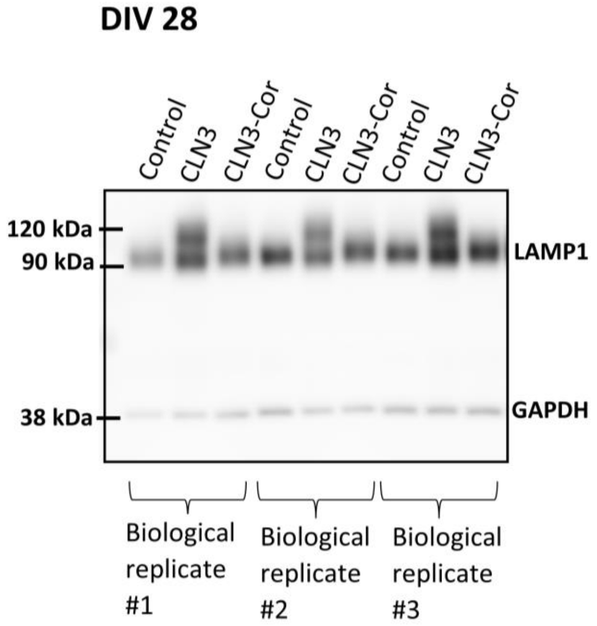

**Fig. 7d**

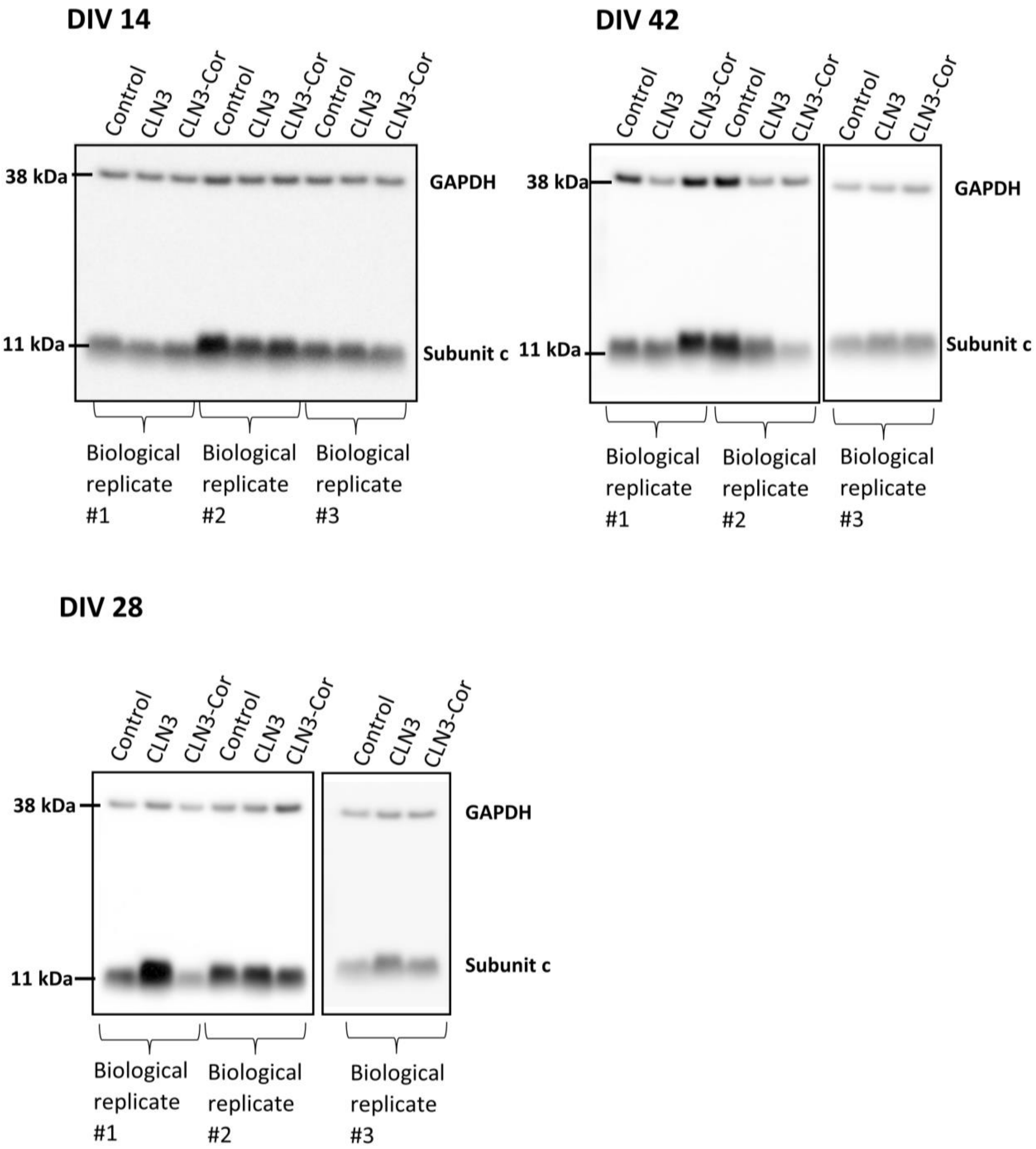
DIV 14
Control
CLN3
CLN3-Cor
Control
CLN3
CLN3-Cor
Control
CLN3
CLN3-Cor
38 kDa
GAPDH
11 kDa
Subunit c
Biological replicate #1
Biological replicate #2
Biological replicate #3
DIV 42
Control
CLN3
CLN3-Cor
Control
CLN3
CLN3-Cor
Control
CLN3
CLN3-Cor
38 kDa
GAPDH
11 kDa
Subunit c
Biological replicate #1
Biological replicate #2
Biological replicate #3
DIV 28
Control
CLN3
CLN3-Cor
Control
CLN3
CLN3-Cor
Control
CLN3
CLN3-Cor
38 kDa
GAPDH
Subunit c
11 kDa
Biological replicate #1
Biological replicate #2
Biological replicate #3

**Figure S7**

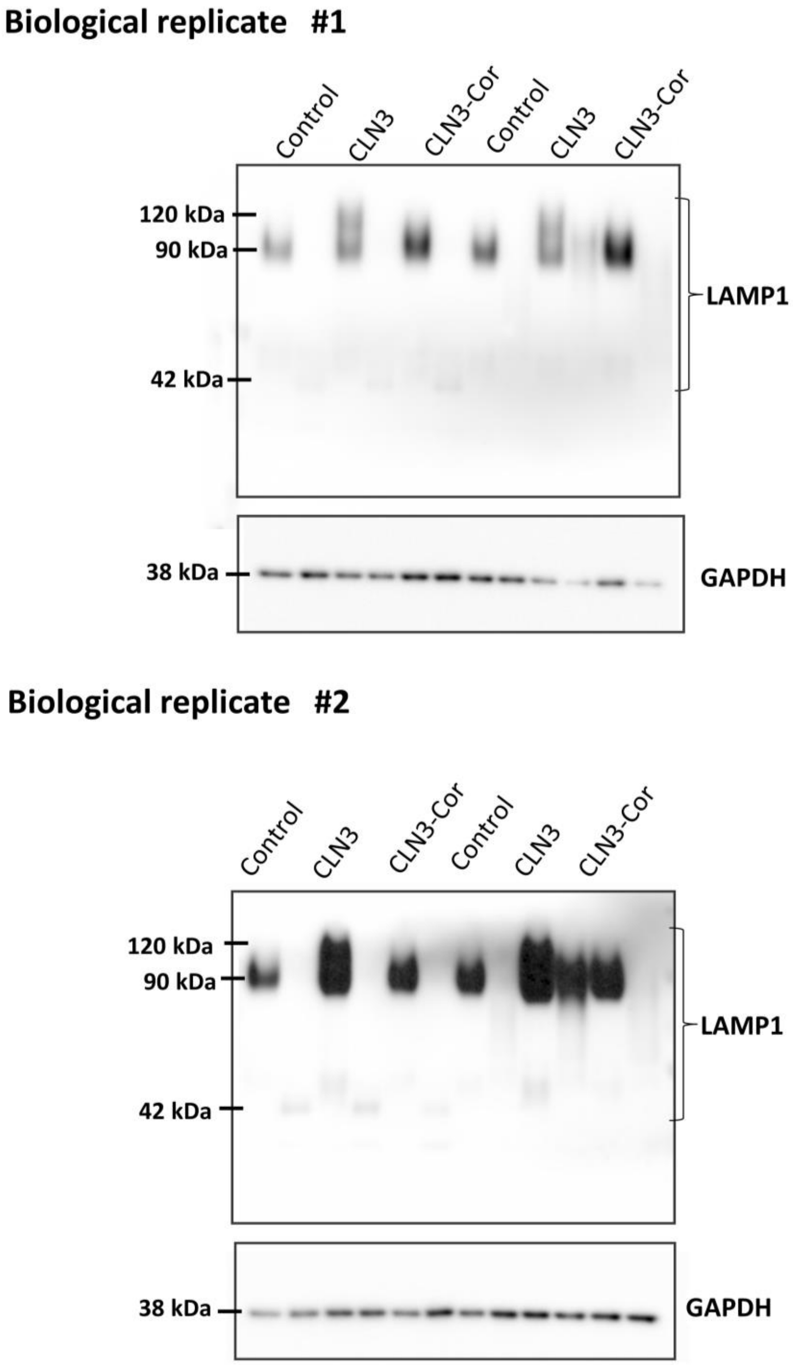
Biological replicate #1
Control
CLN3
CLN3-Cor
Control
CLN3
CLN3-Cor
120 kDa
90 kDa
42 kDa
LAMP1
38 kDa
GAPDH
Biological replicate #2
Control
CLN3
CLN3-Cor
Control
CLN3
CLN3-Cor
120 kDa
90 kDa
42 kDa
LAMP1
38 kDa
GAPDH
